# Age-Related Decline in NCKX4-Mediated Calcium Clearance Accelerates Aortic Remodeling and Drives Early Vascular Aging

**DOI:** 10.1101/2025.10.01.679848

**Authors:** Guilherme H. Souza Bomfim, Kesava Asam, Nish Patel, Kristen Rosenberg, Erna Mitaishvili, Talita Aguiar, Emmanuel Zorn, Ravichandran Ramasamy, Bradley Aouizerat, Rodrigo S. Lacruz

**Author notes:** **Corresponding author**: Guilherme H. Souza Bomfim. Department of Molecular Pathobiology, New York University College of Dentistry, New York, NY, 10010, USA. E-mail address and.

## Abstract

Aging is the primary nonmodifiable risk factor for cardiovascular diseases (CVDs), with older women facing a greater risk of CVDs than age-matched men. Vascular smooth muscle cells (VSMCs) dysfunction and impaired calcium (Ca^2+^) handling are recognized as central contributors to arterial stiffening and calcification. However, the molecular and functional determinants of Ca^2+^ clearance in vascular aging remains a topic of ongoing research. We identify the (<u>N</u>a^+^)-sodium/<u>C</u>a^2+^-calcium (<u>K</u>^+^)-potassium-dependent e<u>x</u>changer <u>4</u> (<u>NCKX4</u>) as a critical functional regulator of VSMCs Ca²⁺ clearance and vascular integrity. We demonstrate that NCKX4 (coded by *Slc24A4*) expression is markedly reduced in aorta of aged (72-78 weeks) mice, with a pronounced decline in females. Functional assays revealed impaired Ca^2+^ clearance in both aged and *Nckx4*^⁻^*^/^*^⁻^ VSMCs, which was accompanied by increased calcification. Histomorphometric analyses of young *Nckx4*^⁻^*^/^*^⁻^ mice revealed fragmentation of elastic fibers, collagen accumulation, wall thickening, and extracellular matrix (ECM) remodeling, all hallmarks of vascular aging that closely resembled those of aged wild-type mice. Transcriptomic profiling of VSMCs showed that loss of NCKX4 alters pathways linked to Ca^2+^-integrin signaling, ECM turnover, and mineralization, including dysregulation of protective anchorage integrins, microfibril-stabilizing, osteogenic drivers and pro-fibrotic integrins. These findings support a model in which impaired Ca^2+^ clearance promotes maladaptive inside-out integrin signaling, disrupting VSMCs anchorage, ECM homeostasis, and mineralization processes. Collectively, our results establish NCKX4 as a previously unrecognized determinant of vascular aging, whose decline accelerates premature arterial remodeling and calcification. This study positions NCKX4 as a potential mechanistic link between age, sex-dependent vulnerability, and vascular stiffening, with implications for novel therapeutic strategies targeting Ca^2+^ handling in CVDs prevention.

## 1. INTRODUCTION

Aging is the most significant risk factor for many chronic metabolic diseases including various types of cardiovascular diseases (CVDs), which remain the leading cause of morbidity and mortality worldwide ^[1, 2]^. CVDs continues to be the leading cause of mortality in the U.S.A, and it is possibly the main reason for the stagnant growth in life expectancy, accounting for ∼930,000 deaths in 2020 ^[1]^. The elderly population faces a heightened vulnerability to CVDs, with an incidence rate of approximately 75% for those aged 60-79 years and around 85% for individuals above age of 80 years ^[3]^. Older women have a higher risk of CVDs than age-matched men, accounting for 1 in every 5 deaths ^[3, 4]^. A hallmark of vascular aging and CVDs in the elderly is the dysregulation of calcium (Ca^2+^) homeostasis in vascular smooth muscle cells (VSMCs) ^[5, 6]^. This Ca^2+^ imbalance drives functional decline of the arterial wall, contraction-dependent vascular tone and contributes to pathological processes including arterial stiffening, calcification, and maladaptive remodeling through alterations in the extracellular matrix (ECM) ^[7, 8]^. Despite these clear associations, current therapies remain largely symptomatic and non-specific, underscoring a significant unmet need for cellular targeted interventions that address the functional and molecular basis of sex- and age-related vascular dysfunction ^[4, 9, 10]^. Dysfunctional regulation in the Ca^2+^ homeostasis not only alters vascular mechanics, but also engages downstream signaling cascades that could exacerbate arterial stiffening and mineral deposition ^[5, 11, 12]^. These processes converge on the vessel wall to accelerate remodeling, thereby recognizing the regulation of Ca^2+^ as a central mechanism that links VSMCs aging to the pathogenesis and high incidence of age-associated CVDs ^[5, 8, 13]^.

VSMCs occupy the majority of the vessel cellular content and wall volume ^[14–16]^, and are associated with a pivotal role in regulating cytosolic Ca^2+^ levels ([Ca^2+^]_cyt_) to maintain vascular signaling and integrity ^[6, 8]^. [Ca^2+^]_cyt_ is critically regulated by a balance between Ca^2+^ clearance (removal) and Ca^2+^ uptake (entry) processes ^[12]^. Although mammalian cells are equipped with a Ca^2+^-tool kit composed by several Ca^2+^ clearance proteins, only the sodium (Na^+^) calcium (Ca^2+^) exchangers (NCKXs/NCXs) have a high Ca^2+^ clearance capacity ^[17–20]^. It is estimated that Na^+^/Ca^2+^ exchangers are 10- to 50-fold more efficient than plasma membrane Ca^2+^-ATPases (PMCAs) ^[19,21–23]^. The Na^+^/Ca^2+^ potassium-(K^+^)-dependent exchanger 4, (NCKX4), is responsible for the Ca^2+^ clearance of large amounts of cytosolic Ca^2+^ per milliseconds ^[17–19]^. NCKX4 is encoded by the *Slc24a4* gene, and uses an additional K^+^ in the Ca^2+^ exchange gradient. This K^+^ outward gradient allows NCKXs to function efficiently under lower Na⁺ gradients or during conditions where intracellular Na⁺ accumulates ^[24–27]^. NCKXs exchangers are 3-10-fold more efficient compared to the NCXs ^[24–27]^. However, unlike members of NCXs family that are well-documented ^[28–32]^, the role of the K^+^-dependent Na^+^/Ca^2+^ exchangers, NCKXs, is poorly understood, particularly in the context of Ca^2+^ signaling of VSMCs and CVDs. Cytosolic Ca^2+^ levels controls VSMCs contractile profile, stiffness, adhesion and senescence factors through mechanisms that depend on integrin-mediated interactions between VSMCs and ECM ^[8, 33]^. Integrins are heterodimeric transmembrane receptors that provide a key physical interconnection between the ECM and cytoskeleton, maintaining the VSMCs homeostasis and their microenvironment ^[7, 34]^. Downstream signals could be bidirectionally transmitted both from inside-out and outside-in mechanisms regulating cell signaling and ECM for remodeling ^[35, 36]^. With aging, disruption of this integrin-ECM crosstalk promotes maladaptive ECM turnover, elastic fiber fragmentation, and collagen accumulation, leading to progressive arterial remodeling ^[34, 37]^. This integrin-dependent imbalance was described as mediated by altered intracellular Ca^2+^ concentration ^[35]^. A report using healthy VSMCs from adult mice showed that changes in intracellular [Ca^2+^]_cyt_ levels affect the integrin activation through inside-out signaling pathways leading to alteration in VSMCs-ECM interaction ^[8]^. Similarly, Ca^2+^ dysregulation engages signaling cascades that directly contribute to Ca²⁺ overload in VSMCs and promotes phenotypic switching toward an osteogenic-like state, characterized by the activation or suppression of vascular calcification drivers ^[8, 38, 39]^.

While the functional role of Ca^2+^ clearance through NCKX4 exchangers in the VSMCs remains unknown, genome-wide association studies (GWAS) ^[40, 41]^ and microRNA (miRNA) ^[42]^ analyses observed a correlation of NCKX4 with vascular calcification and risk of CVDs development. Additionally, molecular analysis using the major *Slc24a4* transcripts detected high levels of NCKX4 in rat aortic tissue ^[23]^. Experimental studies in rodents have been indispensable in dissecting mechanisms related with age-related CVDs ^[9, 10, 43, 44]^. Vascular aging in mice and rats mirrors many features of the human condition, however, age classification in cardiovascular studies is variable ^[45]^. Its well-accepted that mice ranging ∼72-78 weeks-old are widely described with similar human-equivalent ages and disordered physiological processes in humans ^[45–47]^. Recent aging cardiovascular studies considered juvenile (2-month-old), middle-aged (9-2-month-old) and aged (23-24-month-old) mice that corresponds to equivalent human ages of approximately ∼16, ∼42, and ∼69 years, respectively ^[47, 48]^. A report using genetic mouse model of premature aging showed a reduction of NCKX4 in aorta medial layer and VSMCs from klotho-deficient mice (*kl/kl*) ^[49]^. These observations raise the possibility that an impaired NCKX4-mediated Ca²⁺ clearance may represent a previously unrecognized mechanism associated with age-related CVDs mediated by VSMCs from large arteries such aorta. Therefore, the purpose of this study was to analyze the function of the NCKX4 in VSMCs from aortic tissue, and its pathogenesis underlying the development of vascular aging, aortic remodeling and calcification processes. We identify NCKX4 as a critical determinant of VSMCs Ca^2+^ clearance and vascular integrity during aging. We demonstrate that NCKX4 expression is high in young aortic tissue and in isolated VSMCs, but declines with age in both sexes, with a more pronounced reduction in females. Functionally, loss of NCKX4 either through genetic deletion or natural aging results in abnormal Ca²⁺ extrusion, disruption of ECM architecture, and accelerated aortic remodeling. By linking impaired Ca^2+^ clearance to maladaptive integrin-ECM signaling and mineralization pathways, we positioned NCKX4 as a novel driver of vascular aging. These findings suggest that its decline may represent a key molecular event driving aortic stiffening and calcification during aging.

## 2. MATERIALS AND METHODS

### 2.1 Animal Use and Aged Mice

All procedures employed in this study were conducted in accordance with guidelines approved by the Institutional Animal Care and Use Committee (IACUC) of New York University College of Dentistry (protocol # IA16-00625). Experiments were carried out in male and female NCKX4 knock-out (*Nckx4^-/-^*) strain generously provided and developed by Dr. Haiqing Zhao ^[50]^. Wild-type (WT) controls were obtained by inbreeding C57BL/6 mice and confirming the homozygous presence of *Nckx4* alleles (C57BL/6 wild-type^+/+^) in littermates. These mice were considered as WT littermates. All mice used in the study will undergo validation through genotyping, using specific oligonucleotide primers. PCR amplification of genomic DNA with primers (AS67/68) is expected to yield a size of 450 bp for the *Nckx4* knocked-out exon. See **figure S1** for genotyping of mice to detect the WT allele and homozygous *Nckx4* allele deletion (*Nckx4^−/−^*). Aged mice are used for both aging and senescence studies because of their similarity with disordered physiological processes in humans ^[44, 45, 48]^. WT and *Nckx4^-/-^* mice were assigned in young (Y) groups with 12-15 weeks, while mice were considered aged (A) groups ranging between 72-78 weeks, as widely described in recent aging reports ^[45–47]^.

### 2.2 Primary VSMCs Cell Culture

Primary VSMCs were obtained from mouse aorta using enzymatic digestion and subculturing method ^[51, 52]^. Briefly, thoracic aortic tissues from young and aged WT and *Nckx4^−/−^* mice were dissected using fine-tipped forceps to remove adjacent soft and adventitial tissues, and residues using stainless steel disposable scalpels, size 10 (ExcelInt, cat # 29550) in a balanced Phosphate Buffered Saline solution (PBS, Thermo Fisher cat# 10010049). Aorta from 2-3 mice (20-30 mg) were transferred into filter-sterilize solution containing Amphotericin B (Fungizone, Thermo Fisher cat# 15290018) 1:1000 with complete high glucose Dulbecco’s Modified Eagle Medium (DMEM, Thermo Fisher cat# 11995073) containing 10% Fetal Bovine Serum (FBS, Thermo Fisher cat# A5256701) and 1% Penicillin-Streptomycin 10,000 U/mL (P/S, Thermo Fisher cat# 15140122). Under a tissue culture hood using a clean 100 mm petri dish, the aortic tissues were cut into fragments of ∼1-3 mm by forceps and scalpel #10. To obtain clusters of VSMCs populations, aortic fragments were transferred to 5-mL tubes containing 200 μL of Collagenase Type II from *Clostridium histolyticumrom* (Thermo Fisher cat# 17101015) at 1.5 mg/mL (activity 340 units/mL). This was maintained for 5h at 37 °C in a 5%-CO_2_ incubator, manually mixed every 30 min. The enzymatic reaction was stopped by adding 3 mL of complete high glucose DMEM and centrifuged at 300 X g for 5 min. After remove the DMEM, the pellet was washed and resuspended in 600 μL of fresh DMEM media. 24-well plates were used to plate the VSMCs that remained undisturbed for 5 days in the incubator (37 °C in a 5%-CO_2_). Upon reaching ∼85-90% confluence, the clusters of VSMCs were subcultured at passages P2-P3 using 200 μL of Trypsin-EDTA (0.25%, Thermo Fisher cat# 25200056) for 5-7 min to detach the VSMCs. VSMCs were plated on 6-well plates or on glass cover slips coated with Corning™ Cell-Tak (Thermo Fisher cat# CB-40240).

### 2.3 qRT-PCR, Western blot and Immunofluorescence

Real-time qRT-PCR and Western blot analysis were performed as we recently described ^[53]^. Total RNA of the cleaned aortic, brain, heart, stomach and liver tissues was obtained using a phenol/guanidine-based lysis and silica-membrane-based purification method (**Fig. S2**). Tissues were placed in pre-filled bead (1.4 mm) mill tubes (Fisher Scientific cat# 15340153) containing QIAzol Lysis Reagent from RNeasy Mini Kit (Qiagen cat #217004), and processed as indicated by the manufacturer. The samples were homogenized using the 6 Six-position microtube homogenizer (Benchmark Scientific, D1036) for 30s, 2 cycles, 15 off, and speed 7.0. Subsequently, the reverse transcription was performed using the iScript cDNA Synthesis Kit (Bio-Rad cat #1708891). For qRT-PCR, we used the SsoAdvanced Universal SYBR Green Supermix (Bio-Rad cat #1725271), and experiments were performed in a CFX Connect Optics Module thermocycler (Bio-Rad). β-Actin was used as the housekeeping gene. Expression levels were calculated using the delta ΔCT (2^−ΔCT^) method, instead of 2^^−ΔΔCT^ fold-changes, for a general comparation between the levels in young and aged groups. For Western blot analysis, total protein was extracted from aortic tissue using Pierce RIPA Lysis and Extraction Buffer (Thermo Fisher cat #89900) with added 1:100 Halt Protease Inhibitor Cocktail (100x, Thermo Fisher cat #1861277) and 1:1,000 phenylmethylsulfonyl fluoride. Western blot samples were homogenized using the same settings for the qRT-PCR, and then cleared by centrifugation (15,000 g, 15 min, 4°C). Proteins were quantified using Pierce BCA Protein Assay Kit (Thermo Fisher Scientific cat #23227). Protein samples were prepared by gentle heating at 37°C for 20 min and 20 μg of protein was equally loaded onto 10% SDS-PAGE resolving gels. Proteins were then transferred to a PVDF membrane and blocked with 5% milk in TBST for 1 h before loading with primary antibody overnight. The following day, membranes were washed three times for 15 min with TBST and loaded with goat anti-rabbit HRP conjugated secondary antibody for 75 min (Bio-Rad cat #1705046). Membranes were washed three times for 15 min with TBST and imaged with SuperSignal West Pico PLUS Chemiluminescent Substrate (Thermo Fisher cat #34577). For loading, control membranes were stripped and reblocked with 5% milk in TBST for 1 h, and reloaded with a primary antibody overnight. Antibodies against GAPDH 1:1,000 (14C10 rabbit mAb, Cell Signaling Technology cat#2118S) and SLC24A4 (NCKX4) polyclonal 1:500 (Abcam cat #136968) and were used. Uncropped and unprocessed immunoblot membranes of *Slc24a4* (*Nckx4*) and GAPDH for each sample are shown in **Figure S3**.

Immunofluorescence staining was performed in order to detect and measure the NCKX4 expression in aortic tissue and in primary VSMCs, as we previously reported ^[53, 54]^. Formalin-fixed paraffin-embedded aortic tissue blocks were deparaffinized and the Diva Decloaker (Biocare Medical cat# DV2004LX) citrate-based buffer was used to perform antigen retrieval and slides were subsequently blocked with 3% BSA (Thermo Fisher cat # AAJ61655AP) for 1h at room temperature. Slides were subsequently incubated overnight at 4°C with Anti-NCKX4 1:350 monoclonal antibody NeuroMab clone N414/25 (UC Davis/NIH NeuroMab, cat # 75404) followed by 3 washes with PBS (Thermo Fisher cat# 10010049). Next, the slides were incubated for 1h at room temperature with a cross-adsorbed secondary antibody 1:800 Goat anti-Mouse IgG (H+L), Alexa Fluor™ 488 (Thermo Fisher cat# A32723) followed by 3 washes with PBS (Thermo Fisher cat# 10010049). The slides were mounted with Vectashield Vibrance^®^ Antifade Mounting Medium with DAPI (H-1800) (Vector laboratories cat# H-1800-10). Representative images of aortic tissue and primary VSMCs controls from mouse lacking NCKX4 (*Nckx4^−/−^*) are shown in supplementary material (**Fig. S4**). Also, the validation of NCKX4 and specificity of the secondary antibody was performed using only Goat anti-mouse Alexa Fluor 488 (**Fig. S5**).

### 2.4 Measurements of Exchangers Activity and Ca^2+^ Clearance

To measure the Na^+^/Ca^2+^ exchangers activity and the Ca^2+^ clearance capacity of the NCKX4, the NCKXs/NCXs were analyzed in the reverse mode (Ca^2+^ influx) and forward mode (Ca^2+^ clearance) using real-time record of cytosolic Ca^2+^ transients, as we previously described ^[53]^. VSMCs were incubated for 1 h at room temperature with 1 µM of the ratiometric Ca^2+^ probe Fura-2 AM (cat #F1221; Thermo Fisher Scientific) in a regular VSMCs buffer’s solution containing (in mM): 135.0 NaCl; 5.0 KCl; 2.5 CaCl_2_; 1.0 MgCl_2_; 10 D-glucose; and 10 HEPES (pH 7.35-7.45 with NaOH). All reagents and chemicals were obtained from Sigma-Aldrich. Cytosolic Ca^2+^ transients were measured at room temperature (24 ± 2°C) using a polycarbonate 260 μl chamber (cat #RC21BR; Harvard Bioscience Inc.) mounted on an inverted microscope coupled to a perfusion system electrically controlled. Fluorescence recordings were obtained using the Nikon Ti2-E Eclipse inverted light microscope, equipped with an objective (Nikon S Fluor × 20; numerical aperture: 0.75) and a digital SLR camera (DS-Qi2; Nikon) controlled by computer software (NIS Elements version 5.20.01). VSMCs were continuously perfused by a six-way perfusion system (VC-8 valve controller) at 5-6 ml per minute with a common outlet 0.28-mm tube driven by controlled valves (Harvard Bioscience Inc.). The Ca^2+^ probe Fura-2 AM was excited alternatively at 340 and 380 nm using a Lambda LS xenon-arc lamp (Sutter Instrument) and/or pE-340 Fura (Cool Led). Emitted fluorescence was collected through a 510-nm emission filter. All fluorescence images were generated at 5-s intervals, normalized, and the ratio values were calculated using Image J (1.53J).

In order to quantify the Na^+^/Ca^2+^ exchangers activity and Ca^2+^ clearance capacity of NCKX4, the reverse mode (Ca^2+^ influx) and forward mode (Ca^2+^ clearance) of NCKX/NCX were analyzed as previously ^[53]^. The exchangers activity in the reverse mode was elicited by the replacement of Na^+^ (NaCl) with N-methyl-D-glutamine (NMDG, Sigma-Aldrich cat #66930) and maintaining the osmolarity and equimolar amounts of NMDG and NaCl. This was performed in a Ca^2+^-free buffer solution followed by the reintroduction of Ca^2+^ (2.5 mM of CaCl_2_) at 60s. This allows the reversion of Na^+^ and Ca^2+^ electrochemical gradient, increasing the levels of cytosolic Ca^2+^ via NCKXs/NCXs in the reverse mode (Ca^2+^ influx) ^[19, 21, 23]^. Upon reaching Ca^2+^ peak (300s) in the reverse mode, the forward mode of Ca^2+^ clearance was elicited by reintroduction of Na^+^ solution in a Ca^2+^-free solution, returning the Ca^2+^ to basal levels (480s). Additionally, the Ca^2+^ clearance capacity of NCKX4 was also measured after eliciting a high increase of cytosolic Ca^2+^ levels mediated by store-operated Ca^2+^ entry (SOCE) through ORAI channels. SOCE was stimulated by pre-incubation with 1 μM of thapsigargin (Sigma-Aldrich, USA; cat# T9033) for 15 min prior to the experimental protocols in Ca^2+^-free buffer solution. Subsequently, under fluorescence recordings, VSMCs were continuously perfused for 60 s with Ca^2+^-free buffer solution, followed by a re-addition of 2.5 mM of extracellular Ca^2+^in a regular VSMCs buffer’s solution to measure the SOCE peak. At 160s the Ca^2+^ clearance forward mode was elicited by replacement of Ca^2+^ solution with a with Ca^2+^-free buffer solution, returning the Ca^2+^ to basal levels (420s). All the chemical and pharmacological validation of NCKXs/NCXs activity in Ca^2+^ influx reverse mode, effects of Ca^2+^ replacement, and identity of exchangers were detailed in the supplementary material (**Fig. S6**).

### 2.5 Alizarin Red S Staining

VSMCs were maintained in culture for 16 days in a procalcifying DMEM media containing calcium chloride (CaCl_2_, 2 mM, Sigma-Aldrich, USA; cat# 21115) and β-Glycerol phosphate disodium (β-gly, 10 mM, Sigma-Aldrich, USA; cat# 50020), as previously described in VSMCs ^[51]^. Briefly, to measure Ca^2+^ nodule formation, calcification and mineralization, VSMCs were rinsed in sterile PBS twice and fixed with 1 mL of 4% of PFA (Thermo Fisher, cat #J61899.AP) for 15-20 min at 20-25 °C. Both VSMCs exposed to 16-days of procalcifying media and its control cultured in only high glucose regular DMEM were stained with 1 mL of Alizarin Red S 2% (w/v) (Sigma-Aldrich cat# A5533) for 30 min at 20-25 °C. Next, under gentle agitation, and the excess unbound dye removed washing three times with PBS. Images of mineralized nodules were taken with a light microscope. For quantitative analysis of the Alizarin Red S staining, VSMCs were incubated with 1 mL of Hexadecyl Pyridinium 10% (w/v) (Sigma-Aldrich cat# C9002) for 1h at 20-25 °C. ∼400 uL of supernatant was collected to measure the absorbance at 550 nm using a multi-mode microplate reader FlexStation 3 (Molecular Devices, USA) ^[55]^. Negative control images of Alizarin Red S staining in VSMCs exposed to a non-procalcifying medium (high glucose regular DMEM) were showed in the supplementary material (**Fig. S7**).

### 2.6 Aorta Histomorphometric Analysis

Thoracic aorta segments were processed for histomorphometric analysis, as previously described ^[56, 57]^ by the NYU Langone’s Experimental Pathology Research Laboratory (RRID:SCR_017928). Aortic slides were stained with Elastin, Movat’s Pentachrome or Hematoxylin and Eosin (H&E). Elastin staining was used to observe elastic fibers. The Movat’s Pentachrome staining allows to visualize elastic fibers, ECM components, mucin, proteoglycans/ECM, collagen and VSMCs/fibrinogen. H&E staining was used by a blinded researcher to analyze the aortic remodeling measured by histomorphometric parameters of the total wall thickness, tunica media thickness, tunica adventitia thickness and nuclear area. Photomicrographs-images sections were acquired and digitalized using the digital slides Aperio CS2 image capture (Leica Biosystems, USA), 20X magnification lens (UPLXAPO, 0.80NA, 0.6mm WD) coupled to Aperio ImageScope software version 12.4. All histomorphometric parameters were calculated using ImageJ/Fiji (1.54p/J) ^[56]^.

### 2.7 Bulk RNASeq Analysis

RNA from cleaned aortic tissues was isolated using RNeasy Mini Kit (Qiagen cat #217004), as indicated by the manufacturer. RNA quality and RNA sequencing (RNA-seq) were conducted at NYU Langone’s Genome Technology Center (RRID: SCR_017929), as we previously reported ^[57, 58]^. Briefly, aortic tissue from 2 different mice (∼20mg) were combined, totalizing 5 samples (n=5), for each experimental group and RNA quality was assessed on an Agilent 2100 Bioanalyzer (Agilent, USA). All samples were processed in the same time period, following the same protocol in order to limit the batch effects. Next, the RNASeq enrichment libraries were prepared using the low-input RNA sequencing, TruSeq Stranded RNA Library Prep Gold. The libraries were pooled equimolarly into each lane using standard protocols and sequenced by NovaSeq X+ 10B 100 Cycle Flowcell. Raw paired-end sequencing reads from mouse samples were quality controlled using the FASTQC tool (www.bioinformatics.babraham.ac.uk/projects/fastqc/). FASTQ files were QC’d and adapters were removed using Fastp (version 0.24.0), and the quality reports were checked using multiQC (version 1.25.1). The FASTQ files were processed to get the gene counts that were then used to perform Differential Expression Analysis using DESeq2. Pathway analysis was then performed on the DEG using Clusterprofiler. QC’d sequences were mapped to the mouse genome reference (GRCm39 v113) using the STAR (version 2.7.11b). Gene expression was compared using DESeq2 (v.1.32.0), which assumes a negative binomial distribution using the mean and variance estimated from gene counts and p-values calculated using the Wald test. GSEA was performed using GSEA 3.0 (http://www.broadinstitute.org/gsea/). Adjusted p *<* 0.05 and a false discovery rate (FDR) q *<* 0.25 were considered significant. DEGs were selected when presented with an adjusted pvalue *<* 0.05 and a log2FC (fold change) *>* 1 for upregulated genes and *<* -1 for downregulated genes.

The Database for Annotation, Visualization, and Integrated Discovery (DAVID, http://david.abcc.ncifcrf.gov/, version 6.8) was used to perform GO and Kyoto Encyclopedia of Genes and Genomes (KEGG) pathway analyses. Set enrichment analysis was used for the pathway by selecting non-significant differentially expressed genes specified as the “background universe” and accounting for multiple testing using a false discovery rate of q*<* 0.1. Differentially altered pathways were evaluated using the enrich plot package in R to visualize functional enrichment ^[54]^. Principal Component Analysis (PCA) and of bulk RNA-seq data of aortic tissue volcano plot of differential gene expression of bulk RNA-seq data displays the distribution of all analyzed genes based on their differential expression between experimental conditions. See supplementary figures (**Fig.S8** and **S9**).

### 2.8 Data Analyses and Statistics

All data, mathematical analyses and graphs were analyzed and/or generated using the GraphPad Prism software version 10.5.0 (Inc., California, USA), as we previously described ^[53, 59, 60]^. The bioinformatic analysis of RNA-seq data was performed using different pipelines, package in R software and HPC singularity container, as previously described ^[54]^. Data represent the mean ± SEM of 3-5 independent experiments, each individual dot in the histograms represents a sample or region of interest (ROI) from subset of independent cell and tissue image, as detailed on the caption of each figure. Parameters of integrated density and intensity of mean gray values were calculated in each individual ROI from subset of independent images. Representative original traces of Ca^2+^ transients and mathematical parameters were calculated in each individual cell-ROI. The original traces showed the averaged cells, and the number of cells analyzed are shown as individual dots. The kinetics of the Ca^2+^ transients were analyzed by measuring the activity on reverse mode (Ca^2+^ influx) and forward mode (Ca^2+^ clearance) in each individual VSMCs trace and fitted by the nonlinear regression (curve fit) using exponential mode. The “Plateau Followed by One Phase Association” equation has the following function: Y = IF(X < X0, Y0, Y0 + [Plateau-Y0]*{1 − exp[−K*(X − X0)]}) was used to calculate the Ca^2+^ influx reverse mode parameters from 60 s to 290s. At 300s, the parameters for the Ca^2+^ clearance were fitted by “Plateau Followed by One Phase Decay” with the following function Y= IF(X<X0, Y0, Plateau+(Y0-Plateau)*exp(-K*(X-X0))). The delta (Δ) Ca^2+^ peak parameter was calculated for each individual Ca^2+^ transient trace by the subtraction of maximum fluorescence from the basal/minimum fluorescence intensity, expressed by the same unit of Y axis values. The results were analyzed by one-way ANOVA followed by Tukey’s multiple comparison post-hoc test. Significance was accepted at: *P<0.05; **P<0.01; ***P<0.001. ^@^P< 0.05 = aged female group vs. aged male group; ^#^P< WT young group vs. *Nckx4^-/-^* young group; n.s, non-significant, as indicated in each caption figure.

## 3. RESULTS

### 3.1 Age Reduces the NCKX4 Expression in Aortic Tissue and in Primary VSMCs

The expression profile of the five members of *SLC24A* gene family (*SLC24A1-5*) coding for NCKX1-5 determine their functional relevance in various tissue ^[19, 23, 61]^. Prior Northern blot analysis showed highly expressed NCKX4 levels in aortic tissue of adult rats ^[19]^. Our mRNA expression levels showed that NCKX4 is highly expressed in aortic tissue from both young (Y, 12-15 weeks) male and female WT mice, while aorta from aged (A, 72-78 weeks) mice revealed a decrease of ∼70% in male and ∼50% in female (**Fig. 1A**). By contrast, the mRNA levels of *Slc24a4* in tissues that also express this gene including brain, heart, stomach and liver, *Slc24a4* expression was not altered with age (**Fig. S2**). We further determined the expression of NCKX4 in aortic tissue by Western blot analysis, confirming an age-related reduction in the NCKX4 protein levels both in female and male (**Fig. 1B-C**). In addition, immunofluorescence analysis confirmed that NCKX4 is abundant in aortic tissue of WT young mice with a sex- and age-dependent reduction (**Fig. 1D-E**). Next, in primary cultured VSMCs from aorta of aged animals, NCKX4 levels revealed a drastic reduction. This decrease in NCKX4 expression of VSMCs was ∼70% in aged females, while VSMCs of aged males showed a NCKX4 reduction of ∼44% (**Fig. 1F-G**). VSMCs occupy majority of the cellular content, representing ∼50% of the total volume in large arteries such aorta ^[15, 16]^, suggesting that an age-linked decrease in the NCKX4 expression can affect the Ca^2+^ clearance and Ca^2+^ extrusion capacity of VSMCs, particularly in aged females.

**Figure 1.**
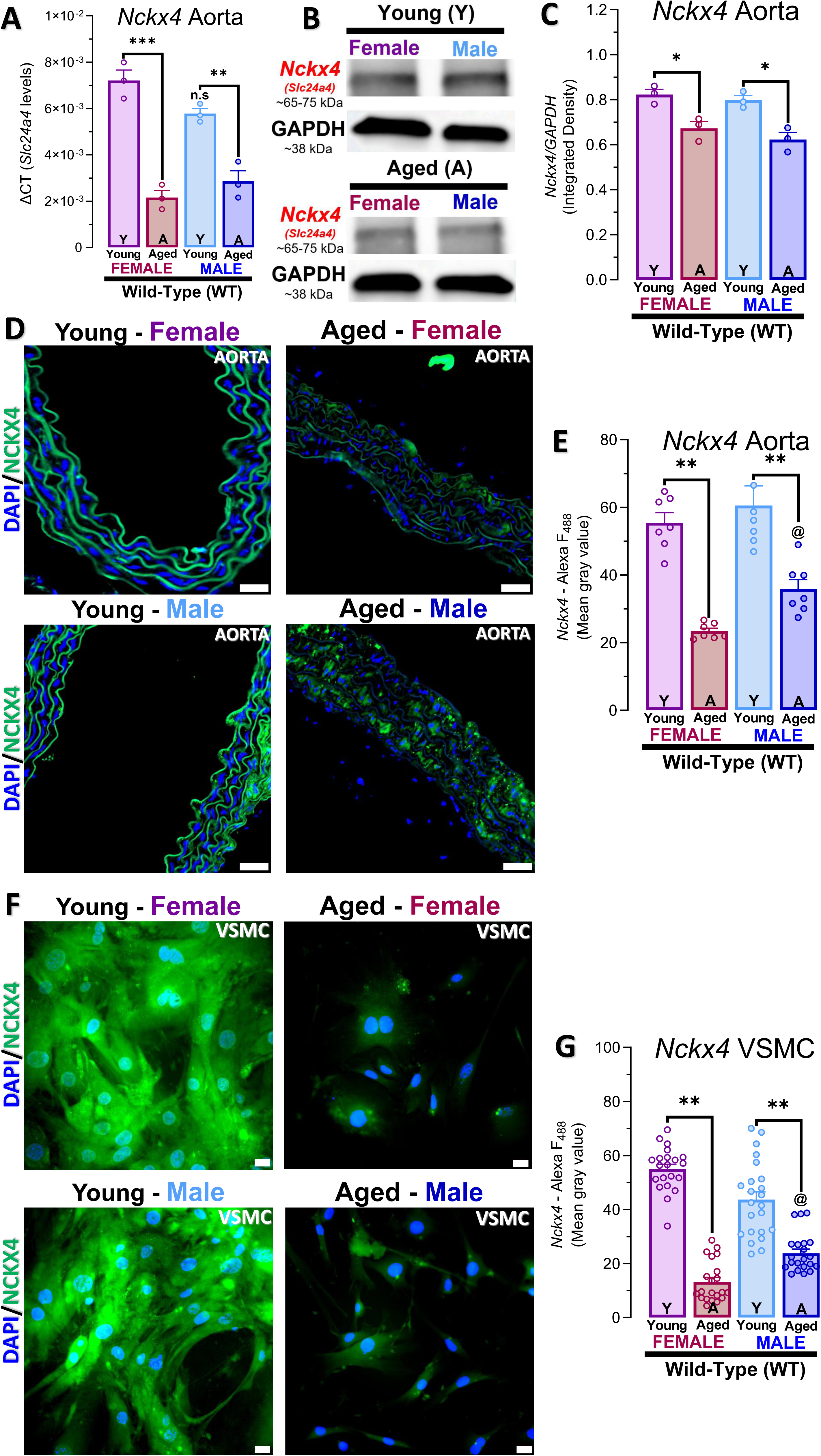
NCKX4 Expression Decrease with Age in Whole Aortic tissue and in isolated VSMCs. Gene expression of *Slc24a4* (*Nckx4*) in thoracic aortic tissue and primary VSMCs from young (Y, 12-15 weeks) and aged (A, 72-78 weeks) male and female wild-type (WT) mice. **(A)**, mRNA levels of *Nckx4* were analyzed by qRT-PCR using the 2^−ΔCT^ method, using β-actin as the housekeeping control. **(B)**, western blot analysis of NCKX4 expression in thoracic aortic tissue predicted NCKX4 band size at ∼65-75 kDa and GAPDH at ∼38 kDa, serving as control. **(C)**, ratio of NCKX4/GAPDH represented as integrated density levels. **(D-E)**, immunofluorescence of NCKX4 in thoracic aortic tissue and in primary VSMCs **(F-G)** from young (Y, 12-15 weeks) and aged (A, 72-78 weeks) male and female wild-type (WT) mice. Anti-NCKX4 is showed in green, and quantified as intensity of mean gray values. Autofluorescent of elastin and elastic fibers at blue excitation light (∼340-400 nm), and nuclear staining of VSMCs with DAPI/Hoechst showed in blue. Data represent the mean ± SEM of ≥3 experiments, and each individual dot in the histograms of immunofluorescence represent an independent region of interest (ROI) from subset of independent images. Data were analyzed by one-way ANOVA followed by Tukey’s multiple comparison post-hoc test. Significance was accepted at: *P<0.05; **P<0.01; ***P<0.001; ^@^P< aged female group vs. aged male group; n.s, non-significant.

### 3.2 Age and *Nckx4^-/-^* Impair the Ca^2+^ Clearance and Na^+^/Ca^2+^ exchangers activity

NCKX4 expression was markedly reduced in aorta and VSMCs of aged (72-78 weeks) female mice. To determine the contribution of age-related decrease on Ca^2+^ clearance and NCKX4 isoform by itself, we measured NCKXs/NCXs activity on reverse mode (Ca^2+^ influx) and forward mode (Ca^2+^ clearance) in VSMCs of female mice. NCKXs/NCXs activity in the reverse mode was elicited by the replacement of external Na^+^ with NMDG, followed by forward mode function elicited by reintroduction of a regular Na^+^ solution (**Fig. 2A-B**). We show that the activity of the Na^+^/Ca^2+^ exchangers on the reverse mode (Ca^2+^ influx) were decreased with age in VSMCs from WT mice. *Nckx4^-/-^* mice showed reduced Na^+^/Ca^2+^ exchangers function, represented by changes in the Ca^2+^ rate and Ca^2+^ peak parameters (**Fig. 2C-D**). The Ca^2+^ clearance in forward mode of Na^+^/Ca^2+^ exchangers showed a drastic reduction in Ca^2+^ extrusion with age. VSMCs from young *Nckx4^-/-^* revealed a reduction in the Ca^2+^ clearance capacity similar to that seen in aged WT mice. The Ca^2+^ clearance rate was reduced by ∼48% in VSMCs of WT aged, ∼30% in young *Nckx4^-/-^*and ∼41% in aged *Nckx4^-/-^* mice (**Fig. 2E-F**). The identity of NCKXs/NCXs was also chemically and pharmacologically confirmed by the use of exchangers antagonist and electrochemical gradient manipulation (**Fig. S6**).

**Figure 2.**
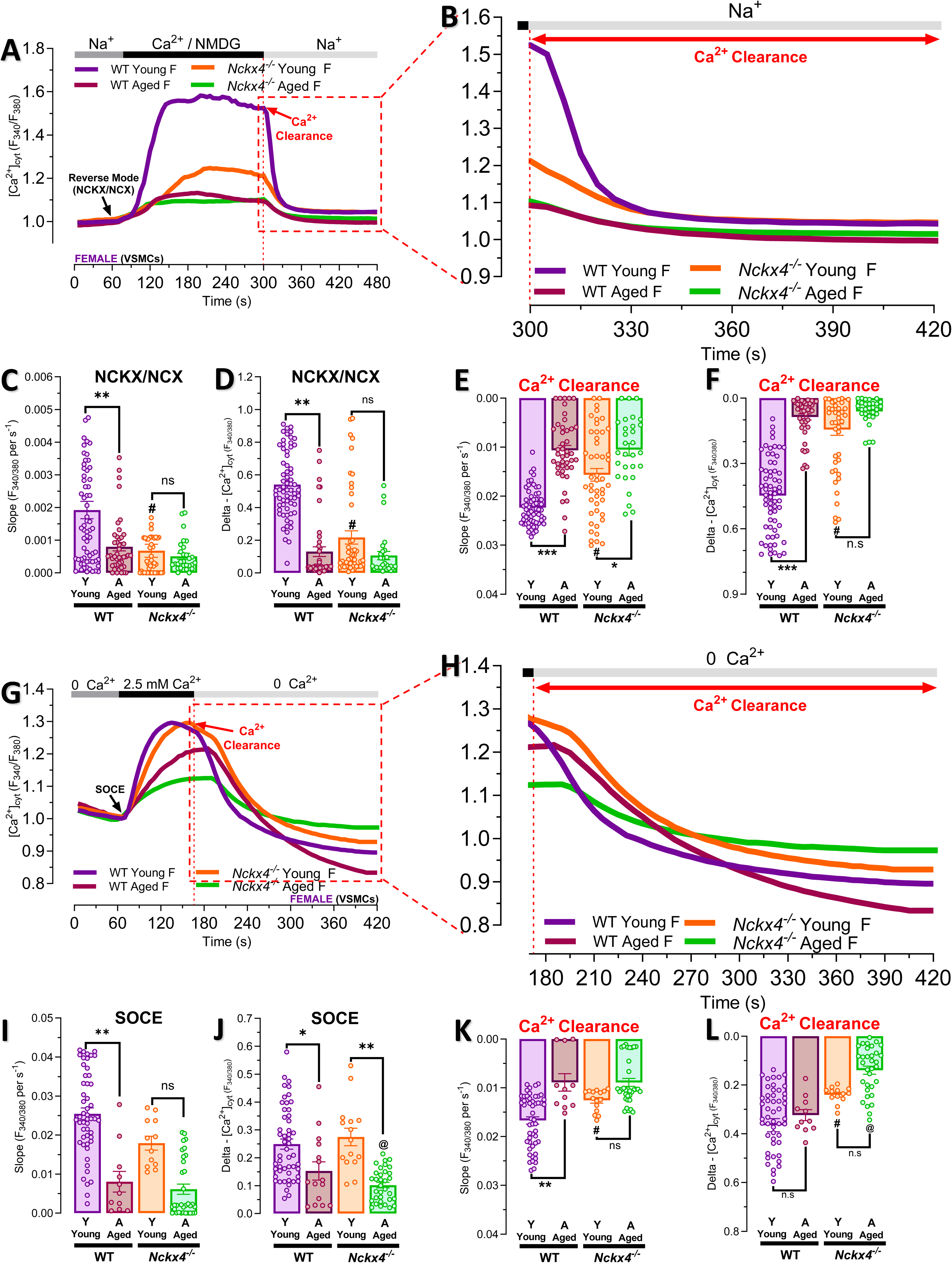
Age and Loss of NCKX4 Impair Ca^2+^ Clearance and Exchangers NCKX/NCX Activity. **(A)**, original traces of NCKX/NCX activity on reverse mode (Ca^2+^ influx) and forward mode (Ca^2+^ clearance) in primary VSMCs from young (Y, 12-15 weeks) and aged (A, 72-78 weeks) female wild-type (WT) and *Nckx4^-/-^* mice loaded with Ca²⁺ probe Fura-2-AM. The NCKX/NCX activity in reverse mode was elicited by replacement of Na^+^ with N-Methyl-D-glucamine (NMDG) and addition of Ca^2+^ (2.5 mM of CaCl_2_) at 60s. **(B)**, upon reaching peak, the forward mode was elicited by reintroduction of NaCl at 300s. Quantification of Ca^2+^ influx rate and Ca^2+^Δ peak parameters in NCKX/NCX activity **(C-D)** and Ca^2+^ clearance **(E-F)**. **(G)**, Ca^2+^ clearance capacity measured after Ca^2+^ influx mediated by SOCE through ORAI channels. SOCE was recorded after pre-incubation with thapsigargin (15 min, 1 μM), followed by perfusion with Ca^2+^-free Ringer solution (60s) before re-addition of 2.5 mM of CaCl_2._ **(H)**, at 160s the forward mode was elicited by reintroduction of NaCl. Quantification of Ca^2+^ influx rate and Ca^2+^ Δ peak parameters of SOCE **(I-J)** and Ca^2+^ clearance **(K-L)**. Data represent the mean ± SEM of ≥3-4 experiments, and each individual dot in the histograms represent a cell labeled as a region of interest (ROI). Data were analyzed by one-way ANOVA followed by Tukey’s multiple comparison post-hoc test. Significance was accepted at: *P<0.05; **P<0.01; ***P<0.001; ^#^P< WT young group vs. *Nckx4^-/-^* young group; ^@^P< WT aged group vs. *Nckx4^-/-^* aged group; n.s, non-significant.

We further investigated the Ca^2+^ clearance capacity of NCKX4 by increasing the cytosolic Ca^2+^ levels via SOCE. At SOCE peak (160s), the Ca^2+^ clearance was measured using a Ca^2+^-free buffer solution with regular Na^+^ concentration (**Fig. 2G-J**). The Ca^2+^ clearance capacity of VSMCs after SOCE induction was reduced by ∼ 24% in young *Nckx4^-/-^*, and ∼46% both in aged WT and aged *Nckx4^-/-^*, represented by Ca^2+^ rate parameter (**Fig. 2K-L**). Taken together, these results suggest that function of NCKXs/NCXs exchangers is impaired with age. Additionally, the early loss of NCKX4 (*Nckx4^-/-^*) alone in VSMCs of young female mice compromises the high Ca^2+^ capacity of NCKXs/NCXs exchangers.

### 3.3 *Nckx4^-/-^* Enhances Calcification and Higher Mineralization Potential

Given that age and *Nckx4^-/-^* impairs Ca^2+^ clearance, we next investigated whether this would affect the capacity of VSMCs to form mineral Ca^2+^ phosphate nodules. Calcification was elicited maintaining VSMCs in a procalcifying DMEM media supplemented with Ca^2+^ (CaCl_2_) and phosphate (β-gly) to induce Ca^2+^ phosphate nodules formation, then stained with Alizarin Red-S and dissolved by hexadecyl pyridinium to measure and quantify the mineral Ca^2+^ phosphate deposits (**Fig. 3A**). As a positive control, VSMCs from both young and aged mice formed mineral precipitates after 16 days in procalcifying media (**Fig. 3B-C**). We show in young *Nckx4^-/-^* enhanced calcification by ∼40% to the similar levels (∼31%) observed in WT aged mice (**Fig. 3C-D**). In addition to age, the NCKX4 absence increased calcification by ∼32% in aged *Nckx4^-/-^*mice (**Fig. 3E-F**). These results suggest that lack of NCKX4 drives an early calcification in young mice, and enhances mineralization in an age-dependent manner. Negative controls of non-mineral Ca^2+^ phosphate nodules formation was performed culturing VSMCs in a non-procalcifying media (**Fig. S7**).

**Fig. 3.**
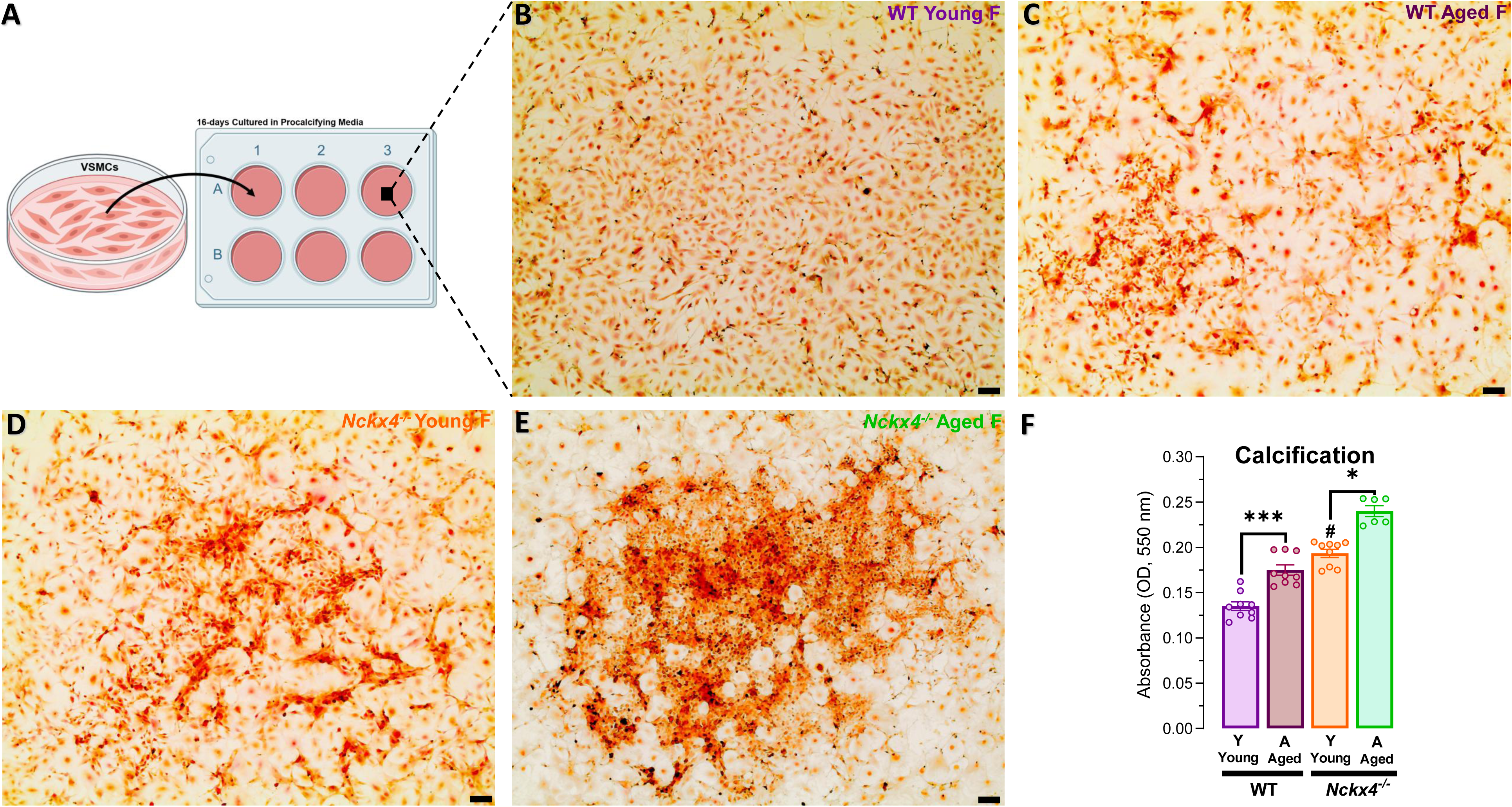
VSMCs from *Nckx4^-/-^* Shown Increased calcification and Higher Mineralization Potential. **(A-E)**, schematic and representative images of mineralized Ca^2+^ phosphate nodules after 16-days of VSMCs calcification using a procalcifying DMEM media containing CaCl_2_ (2 mM) and β-gly (10 mM). VSMCs from young (Y, 12-15 weeks) and aged (A, 72-78 weeks) female wild-type (WT) and *Nckx4^-/-^* mice loaded were stained with Alizarin Red-S (2%, w/v), and dissolved by hexadecyl pyridinium (10%, w/v). **(F)**, mineral Ca^2+^-phosphate deposits were quantified by absorbance of violet destained solution at 538 nm. Data represent the mean ± SEM of ≥3 experiments, and each individual dot in the histograms represent an independent sample from 6-well-plate stained with Alizarin Red-S. Data were analyzed by one-way ANOVA followed by Tukey’s multiple comparison post-hoc test. Significance was accepted at: *P<0.05; ***P<0.001; ^#^P< WT young group vs. *Nckx4^-/-^* young group.

### 3.4 *Nckx4^-/-^* Mice Show Early Aortic Remodeling Consistent with Aged Features

Histological cross-sections from aorta of young *Nckx4^-/-^*female mice revealed marked structural alterations compared to young WT controls. These changes were represented by the loss of pattern of elastic fibers and increased ECM composition with disruption on elastic fibers and ECM remodeling (**Fig. 4**). Elastin staining showed a reduction of the characteristic zig-zag pattern of elastic lamellae, and loss of organized elastic fiber architecture in aortic tissue of young *Nckx4^-/-^* that impair in aged *Nckx4^-/-^* mice (**Fig. 4A-B**). Similarly, Movat’s Pentachrome staining suggests an ECM remodeling associated with increased collagen deposition in aorta of young *Nckx4^-/-^* mice (**Fig. 4C-D**). In aortic cross-sections, Movat’s staining distinctly visualized elastic fibers (black), collagen (yellow), VSMCs/fibrinogen (red), and mucin/proteoglycans (green). Young *Nckx4^-/-^*mice displayed a shift in wall composition with an expansion of green- and yellow-stained regions indicating collagen deposition within the medial and adventitial layers. Additionally, the black-stained elastic fibers appeared disrupted and fragmented, consistent with loss of the normal zig-zag architecture. These qualitative alterations associated with the ECM remodeling phenotype were also observed in aged WT mice, reinforcing that NCKX4 deficiency drives premature vascular structural changes remodeling. Therefore, to further quantify and dissect and the aortic remodeling, we performed histomorphological analysis of H&E staining cross-sections which revealed marked structural alterations in aorta of young *Nckx4^-/-^*mice compared to age-matched WT controls (**Fig. 4E**). When compared to young WT control, aortic tissues from young *Nckx4^-/-^* demonstrated a significant increase in total wall thickness (∼28%), tunica media thickness (∼44%), and nuclear area (∼34%) parameters (**Fig. 4F-I**). These changes indicate cellular and extracellular expansion consistent with hypertrophic vascular remodeling. The medial layer exhibited an increase in thickness, expanded and enriched ECM components, suggesting active early aortic remodeling and structural reorganization.

**Figure 4.**
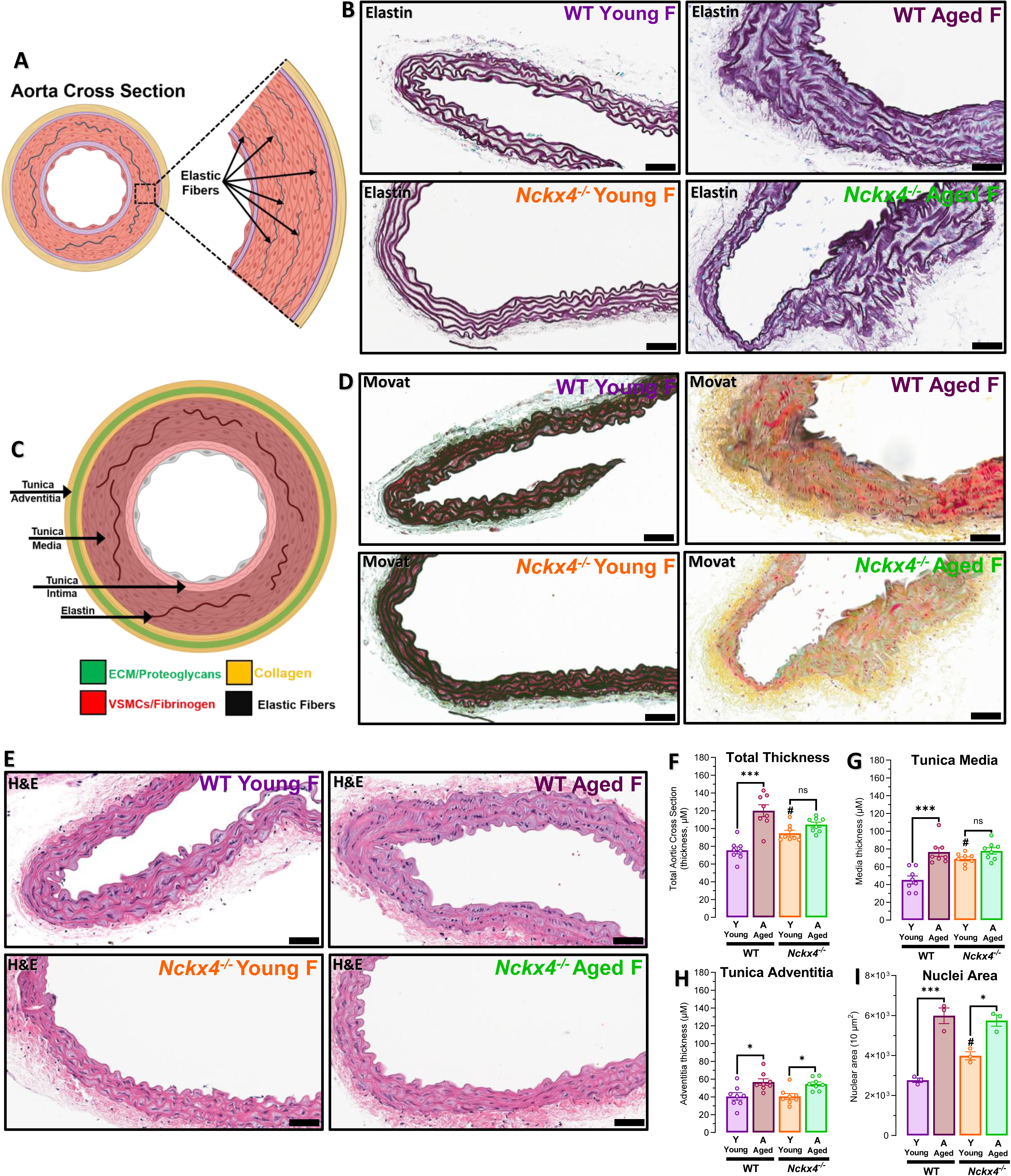
*Nckx4^-/-^* Induces Hypertrophic Aortic Remodeling Consistent with Aging. Photomicrographs and schematics of histological differences in aortic wall of young (Y, 12-15 weeks) and aged (A, 72-78 weeks) female wild-type (WT) and *Nckx4^-/-^* mice. Thoracic aortic tissues were stained with Elastin **(A-B)**, Movat’s Pentachrome **(C-D)** and Hematoxylin and Eosin (H&E) **(E)**. Aortic remodeling was measured by histomorphometric analysis of total wall thickness **(F)**, tunica media thickness **(G)**, tunica adventitia **(H)** and nuclear area **(I)** parameters. Data represent the mean ± SEM of ≥3 experiments, and each individual dot in histograms represent a region of interest (ROI) from 3 different images. Data were analyzed by one-way ANOVA followed by Tukey’s multiple comparison post-hoc test. Significance was accepted at: *P<0.05; ***P<0.001; ^#^P< WT young group vs. *Nckx4^-/-^* young group; n.s, non-significant. Movat’s Pentachrome to visualize elastic fibers, wall and ECM components. Mucin and proteoglycans/ECM = green; Collagen = yellow; VSMCs/fibrinogen = red; Elastic fibers = black. Scale: 20x magnification.

### 3.5 Loss of NCKX4 Causes Early Alteration in VSMCs/ECM via Ca^2+^-Integrin Related Pathways

Because young *Nckx4^-/-^* mice showed compromised Ca^2+^ clearance capacity and induced an aortic remodeling, consistent with aged phenotype features, we used RNA-seq to analyze global patterns of pathways and gene expression changes involved on this phenotype in aortic tissue. PCA and differential expression analysis, showed separated clusters between all experimental groups (**Fig. S8**). As an expected control, RNA-seq analyses identified 1772 differentially expressed genes when WT young was compared with its age-matched WT (**Fig. S9A**). The aorta of young *Nckx4^-/-^*mice showed changes in 1159 genes when compared with the WT aged group (**Fig. S9B**). Also, the lack of NCKX4 altered 636 genes in aged aorta compared with its age-matched WT (**Fig. S9C**). These results indicate that *Nckx4^-/-^*drives changes in the patterns of pathways and gene expression, particularly in aorta of young mice. Next, to further determine whether NCKX4 is involved in vascular aging and in VSMCs-ECM interaction mediated by Ca^2+^-integrin related pathways, we investigated pathways and gene encoding factors associated with ECM remodeling, VSMCs senescence, Ca^2+^-integrin regulation and mineralization. Differential expression analysis identified pathways related to ECM, VSMCs, aortic aging and mineralization (**Fig. 5-6** and **S10**). Although these genes were expected to be altered in aged mice, the lack of NCKX4, mainly in young mice, caused changes on these pathways. In pathways related to collagen processing and ECM degradation, the loss of NCKX4 affected these pathways only in aorta of young animals (**Fig. 5A**). Similarly, pathways related with VSMCs, longevity and Ca^2+^ signaling regulation showed alteration mainly in young mice (**Fig. 5C**). Thus, given the fact that our results suggested an early calcification in VSMCs from young *Nckx4^-/-^* mice, associated with higher mineralization process in age-dependent manner (**Fig. 3**), the pathway analyses identified different Ca^2+^-integrin and calcification related pathways affected by NCKX4 removal in young mice. Loss of NCKX4 in aorta of young animals not only altered pathways related with calcification, Ca^2+^ transport, bone mineralization, aorta development or Ca^2+^-integrin, but also affected the number of genes (count) involved on these pathways, similar to the number found in WT aged group (**Fig. 5-6**).

**Figure 5.**
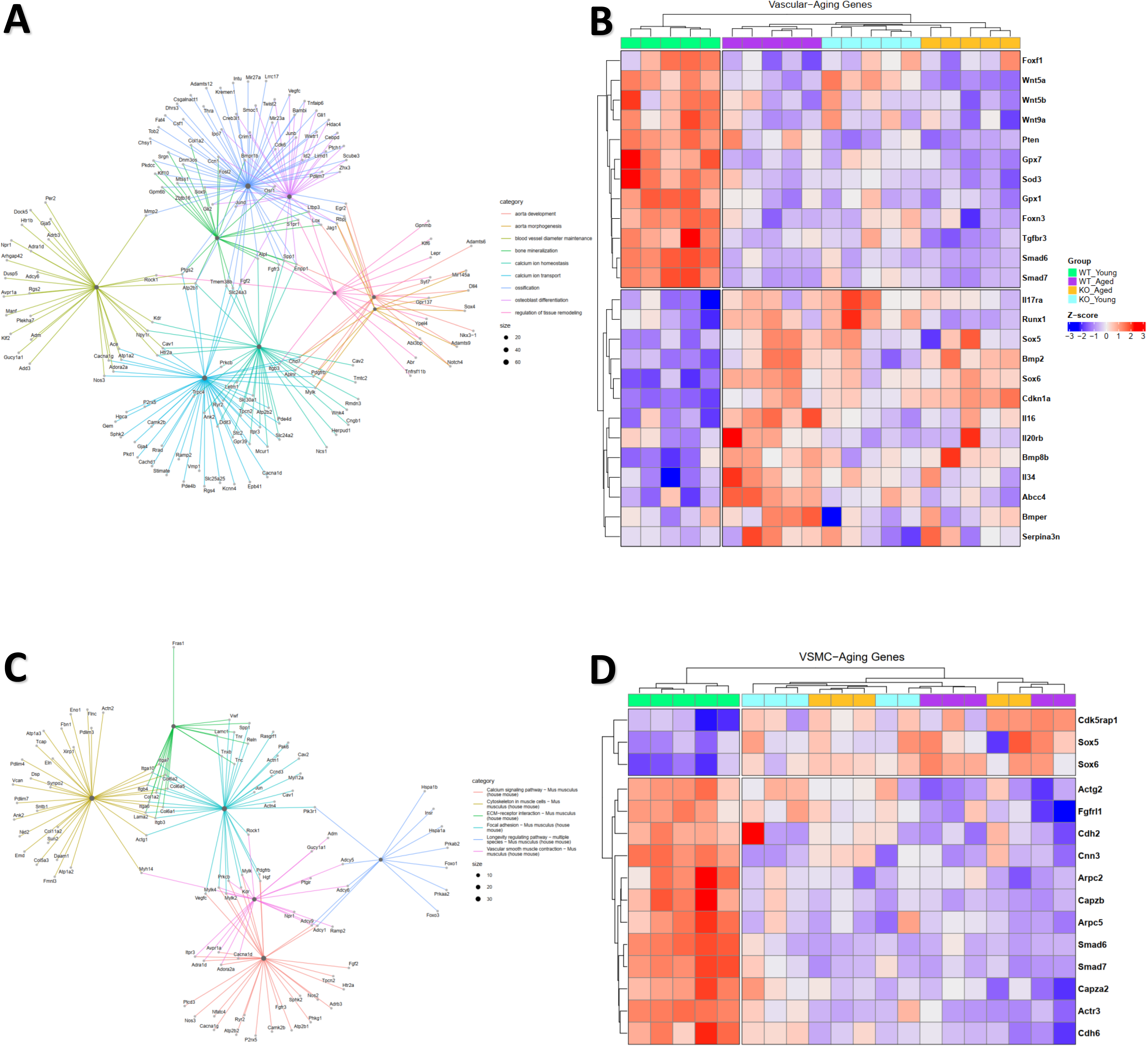
Lack of NCKX4 Drives Early Changes in Vascular and VSMCs genes. **(A,C)**, differential expression analysis for pathways identified genes with significantly different expression levels between experimental conditions. Over Representation Analysis (ORA) targeted pathways related to ECM, VSMCs, aortic aging, Ca^2+^-mediated mineralization. Heatmap representation of vascular aging markers **(B)** and VSMCs genes **(D)**. Normalized values were scaled to convert to z-scores at p< 0.05 with log2FC L1. Individual genes represented as Log2FC changes. Data represent the mean ± SEM of 5 experiments, significance was accepted at: *P<0.05 WT aged group vs WT young group; n.s, non-significant = WT aged group *Nckx4^-/-^* young group or *Nckx4^-/-^* aged group.

**Figure 6.**
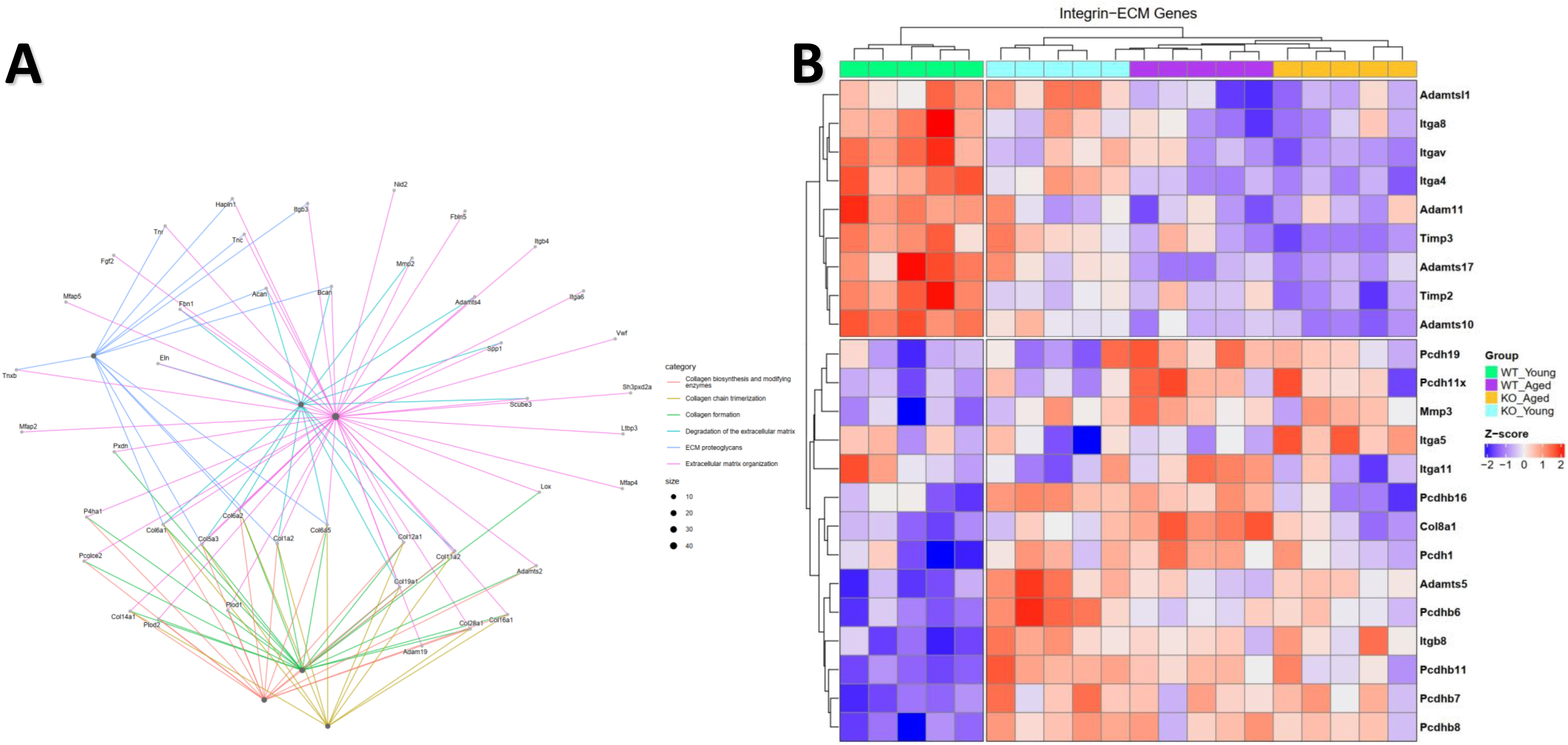
*Nckx4^-/-^* Causes Early Alterations in VSMCs/ECM via Ca^2+^-Integrin Related Pathways. **(A)**, differential expression analysis for pathways identified genes with significantly different expression levels between experimental conditions. Over Representation Analysis (ORA) targeted pathways related to ECM, VSMCs, and integrins. Heatmap representation of integrin mediated VSMCs-ECM interaction genes **(B)**. Normalized values were scaled to convert to z-scores at p< 0.05 with log2FC L1. Individual genes represented as Log2FC changes. Data represent the mean ± SEM of 5 experiments, significance was accepted at: *P<0.05 WT aged group vs WT young group; n.s, non-significant = WT aged group *Nckx4^-/-^* young group or *Nckx4^-/-^*aged group.

Heatmap representation of the differential gene expression profile confirmed that markers of vascular aging were altered in aged WT when compared to the young WT controls. Young *Nckx4^-/-^* showed similar alteration pattern to the WT aged group, followed by non-additional changes in aorta from aged *Nckx4^-/-^* mice (**Fig. 5B**). The young *Nckx4^-/-^*group also revealed alterations consistent with an aged profile of genes associated with VSMCs aging, compared with WT aged group (**Fig. 5D**). Thus, these alterations in gene profile associated with Ca^2+^-integrin related pathways, ECM remodeling and calcification, suggest an involvement of these pathways in early aortic remodeling mediated by NCKX4. Integrin-ECM related genes showed a significant shift in the expression that was both NCKX4- and age-dependent, being consistent with an accelerated vascular aging when Ca^2+^ clearance was impaired by NCKX4 loss (**Fig. 6A**). Genes that encode firm integrin anchorage (*Itga8*, *Itgav* and I*tga4*), ECM microfibril organization (*Adamtsl1* and *Adamts10/17*) and balanced proteolysis (*Timp2/3*) showed high expression in aorta of WT young control mice. However, lack of NCKX4 in young animals or ge caused a progressive downregulation of levels of these genes (**Fig. 6B**). In young WT control mice, we observed low levels of genes related with ECM degradation (*Adamts5* and *Mmp3*), fibrotic integrin-ECM deposition (*Col8a1*, *Itga5/11* and *Itgb8*) and dysfunctional protocadherins (*Pcdh1/19*, *Pcdh11x* and *Pcdhb6/7/8/11/16*). On the other hand, in young *Nckx4^-/-^,* aged WT and aged *Nckx4^-/-^*these genes showed a similar upregulation (**Fig. 6B**). Taken together, the transcriptomic profile suggests that loss of NCKX4, mainly in the aorta of young mice, mediates VSMCs dysfunction, impaired integrin signaling, and alteration in ECM turnover and its properties, promoting early aortic stiffening and calcification that are hallmarks of age-related aortic remodeling.

## 4. DISCUSSION

Aging is the most important nonmodifiable risk factor for CVDs development, and dysfunctional Ca^2+^ handling in VSMCs is a key factor that increases the incidence of CVDs ^[5, 6]^ although the targets, functional and molecular mechanisms have not been fully elucidated. The elderly population faces a heightened vulnerability to CVDs, with an incidence rate of ∼75% for those aged 60-79 years and ∼85% for individuals above age of 80 years ^[3]^. Clinical studies report that older females have a higher risk of CVDs than age-matched men, accounting for 1 in 5 deaths ^[3, 4]^. Despite the observed decline in the function of large arteries with age, there exists an unmet medical need for more targeted therapies ^[9]^. Previous studies both in human and murine VSMCs have shown that aging leads to defective Ca^2+^ handling, indicating the its pivotal role in contraction-dependent aortic stiffness, calcification and remodeling of VSMCs ^[5, 62–64]^. However, the mechanisms that predispose the aging population to dysfunctional Ca^2+^ remain unknown.

Despite the function and mechanism by which NCKX4 drives aged-related CVDs remains unknown, GWAS and miRNAs studies have correlated NCKX4 with potential CVDs and calcification processes ^[40-42]^. A GWAS study in 54 years-old patients (∼58% women) predicted *SLC24A4* as potential candidate gene for systolic blood pressure (SBP) regulation and a future target for pharmacological intervention to reduce the risk of CVDs ^[40]^. The rs11160059, an intronic SNP in *SLC24A4*, showed significant association with SBP among normotensive individuals ^[40]^. By contrast, an independent GWAS using a sample of 2474 unrelated 45 years-old individuals (∼54% women) observed no statistically significant differences in SNP rs11160059 associated with *SLC24A4* ^[41]^. In addition, a murine study using Klotho-deficient mice (*kl/kl*), a genetic model of premature aging, showed that mRNA and protein expression levels of NCKX4 decrease in aortic medial layer of *kl/kl* mice and in cultured VSMCs exposed to high concentrations of extracellular Ca^2+^-phosphate ^[65]^. Aortic VSMCs from ∼2-month-old WT mice overexpressing miR-712* caused a reduction in the mRNA and protein levels of NCKX4 ^[42]^. Both these humans and mice outcomes suggest an association between NCKX4 genetic variants, SBP regulation and mRNA driven genetic modulation. These studies were largely correlative and did not directly investigate the functional role of NCKX4 in aortic VSMCs or how its age- and sex-dependent downregulation contributes to impaired integrin-ECM signaling, Ca^2+^ clearance and aortic remodeling. In this study, we determined that NCKX4 was highly expressed both in aortic tissue and primary VSMCs from young (12-15 weeks) WT mice, with a significant reduction in the aorta of aged (72-78 weeks) (**Fig. 1**). A similar NCKX4 abundance had been previously reported in aorta of young rats with a slight decrease in its levels in aorta of ∼5-month-old male rats ^[23]^. Here we showed that NCKX4 was downregulated particularly in primary VSMCs in a sex- and age-dependent manner (**Fig. 1F-G**). NCKX4 reduction was substantial in both sexes, but more pronounced in females compared with aged males, which may be associated with estrogen and other sex hormones known to regulate Ca^2+^ signaling pathways ^[66–69]^.

Studies using murine models showed that loss of estrogen mediated by ovariectomy or aging causes arterial stiffness, calcification, changes in VSMCs contractile genes, and increase risk CVDs ^[66–68]^. Epidemiologic and clinical reports showed that post-menopausal women have increased arterial stiffness, coronary artery Ca^2+^ and arterial calcification when compared with pre-menopausal women and age-matched men ^[69–71]^. This positions the hormonal deficiency and regulation of Ca²⁺ homeostasis as important drivers for a sex-dependent vulnerability to vascular aging mediated via VSMCs dysfunction.

Throughout the entire vessel wall, VSMCs represent the main cellular content in large arteries like the aorta ^[14]^. In resistance arteries, the proportion of VSMCs to total wall volume is higher, accounting for ∼60-80% of the wall volume ^[14, 16]^, positioning VSMCs to have outsized influence on vessel function. Our findings showed that both the genetic engineered knockdown (*Nckx4^−/−^*) and age-dependent loss of NCKX4 in female cultured VSMCs compromise the high Ca^2+^ clearance and activity capacity of the Na^+^/Ca^2+^ exchangers (**Fig. 2**), suggesting a substantial decline in the capacity of cytosolic Ca^2+^ clearance. In mammalian cells such as VSMCs, the [Ca^2+^]_cyt_ levels are maintained through a sophisticated Ca^2+^ clearance systems, including plasma membrane PMCAs, H^+^/Ca^2+^ transporters, mitochondrial Ca^2+^ uptake and sarco(endo)plasmic reticulum Ca^2+^-ATPases (SERCAs) ^[12, 72, 73]^. As mentioned, both Na^+^/Ca^2+^ exchangers (NCKXs/NCXs) families are responsible for the removal of large amount of cytosolic Ca^2+^, but can also function bidirectionally ^[17–20]^, being approximately 10- to 50-fold more efficient than PMCAs in clearance Ca^2+ [19, 21–23]^. NCKX4 removes 1Ca^2+^ plus 1K^+^ in exchange of 4Na^+^ uptake, with additional use of the K⁺ gradient allowing NCKXs to function 3-10-fold more efficient in remove Ca^2+^ compared to the NCXs exchangers to decrease the levels of cytosolic Ca^2+ [24–27]^.

Huang et al. reported that changes in the cytosolic [Ca^2+^]_cyt_ levels affect integrin activation through inside-out signaling pathways leading to alteration in VSMCs-ECM interaction ^[38]^. The rise in the [Ca^2+^]_cyt_ levels caused a rapid transient increase in VSMCs stiffness, adhesion and contractile event, while the presence of the fast Ca^2+^ chelator, BAPTA, decreased VSMCs adhesion and has no effect on cell stiffness ^[38]^. This suggests a key role of [Ca^2+^]_cyt_ levels in the VSMCs and ECM signaling mediated by Ca^2+^-dependent integrins. In addition to the slow Ca^2+^ clearance capacity of VSMCs from aged and *Nckx4^−/−^*mice (**Fig. 2**), our results indicate that loss of NCKX4 triggers an early calcification in the aorta of young mice, increasing the formation of Ca^2+^-phosphate nodules formation in an age-dependent manner (**Fig. 3**). VSMCs explanted from human aortic tissue revealed a higher mineralization process when cultured in elevated concentration-dependent extracellular Ca^2+^ phosphate levels ^[74]^. Similarly, non-selective pharmacological inhibition of Na^+^/Ca^2+^ exchangers increased the Ca^2+^ content of ECM in human umbilical artery smooth muscle cells ^[74]^. Although the molecular mechanisms remain unknown, the presence of Ca^2+^ chelator BAPTA inhibited the calcification *in vitro* but failed to induce changes in mineralization ^[74]^. A recent report using a model of premature aging found that *kl/kl* mice developed severe arterial calcification and elastin fragmentation ^[49]^. Also, *kl/kl* mice demonstrated higher levels of osteoblast and bone formation makers in aorta, independent of phosphate levels ^[65]^. In this context, translational reports in distal abdominal aorta from aged (60-year-old) female patients showed focal calcification and calcified matrix vesicles, associated with elastin and collagen fibrils, and heavily calcified vessels with mineralization process extended into the ECM ^[74]^. This suggest that the increase in the [Ca^2+^]_cyt_ levels drives an inside-out signals to activate ECM remodeling.

The central underlying pathways related with the risk of CVDs development are linked with vascular aging and associated arterial stiffness remodeling ^[10, 37, 75]^. Arterial remodeling is an independent risk factor for CVDs characterized by structural and functional alterations in tunica media, calcification, and functionality of the VSMCs ^[75–77]^. Large vessels such as the aorta, and medium-sized arteries react in a unique manner to stimuli that induces CVDs when compared to capillaries ^[76, 78]^. A stereotypical vascular injury response is the ECM remodeling that occurs particularly in larger vessels in response to injurious stimuli, such as Ca^2+^ overload, genetic deficiencies and dysfunctional VSMCs-ECM interactions mediated by integrins ^[7, 14, 15, 30]^. Our histological and morphometric analyses demonstrate that young *Nckx4^−/−^*mice develop structural aortic changes that closely resemble those observed in aged WT animals, indicating that NCKX4 deficiency drives premature vascular remodeling (**Fig. 4**). Loss of the organized zig-zag architecture of elastic fibers, fragmentation, and increased collagen deposition within the media and adventitia are hallmarks of age-associated vascular remodeling in both animal models and humans ^[5, 76, 79]^. The expansion of wall and medial thickness is consistent with hypertrophic remodeling driven by ECM and VSMCs reorganization ^[75, 80, 81]^. These changes are active processes that involve VSMCs-ECM interactions, Ca^2+^ signaling, and maladaptive integrin pathways.

Our findings also showed that young *Nckx4^−/−^* mice exhibited elastin alteration and collagen enrichment. Elastic fiber fragmentation and excessive collagen deposition are interconnected with alteration in the expression profile of MMPs and VSMCs-ECM integrin-mediated genes ^[76, 81, 82]^. Elastin-deficient (*Eln^+/−^*) models revealed that elastin insufficiency enhances age-related arterial stiffening and demonstrates the critical role of elastin/collagen ratio in rodent arteries ^[79]^. Females lacking elastin have lower elastin contents in ascending aorta, and smaller outer diameters when compared to age-matched males ^[79]^, suggesting an important sex-dependent differential features during age-related arterial remodeling. A recent study using C57Bl/6 mice revealed a marked age aortic ECM changes, including increased collagen deposits, wall thickness and vessel pressure, and decreased elastin content, which combined represent a hallmark ECM response to aging ^[47]^. The aortic remodeling was age-dependent with alterations ranging from 6-8 to 24 months, but different patterns were observed during these aging stages ^[47]^. We further observed an early aortic remodeling in young (∼3-4-month-old) female mice lacking NCKX4 represented by loss of the architecture of elastic fibers, ECM (green staining) and collagen deposits (yellow staining), associated with wall hypertrophic remodeling (**Fig.4**).

Collagen types I and III are mainly involved in imparting strength to the vessel wall ^[47]^. Additionally, De Moudt et al. reported that collagen type I showed an age-dependent (6 to 24-mounth-old) increase, whereas collagen type III was increased only in the aorta of 24-month-old mice ^[47]^. Other remodeling parameters that supports an early aortic remodeling triggered by the lack of NCKX4 also showed alteration in aorta of WT C57Bl/6 mice with ∼6-8 month, being impaired in age-dependent manner by 24 month ^[47]^. These studies strengthen the notion that impaired Ca²⁺ clearance via NCKX4 deficiency precipitates premature ECM remodeling and mimics phenotypes normally restricted to older individuals. Different human studies in aortic tissues corroborate our findings, with elastin showing pronounced loss and collagen increasing, particularly after mid-life ^[79, 83–85]^, which aligns with the elastic fiber fragmentation and collagen enrichment that we observe in young *Nckx4^−/−^*. In this context, imaging studies in post-menopausal women further show that increased aortic stiffness and calcification are associated with ECM alterations and reduced elastic recoil _[86-88]_. Taken together, our results establish that NCKX4 loss drives early vascular remodeling consistent with aged features, thereby positioning NCKX4 as a pivotal regulator of ECM homeostasis and arterial stiffness. Building on this foundation, the next critical aspect to address is how these ECM and structural changes interface with calcification processes, which further exacerbate vascular remodeling.

To further investigate the mechanisms underlying NCKX4-mediated vascular aging, we performed transcriptomic analyses of the aorta that converged on the notion that loss of NCKX4 accelerates vascular aging by alteration of targeted pathways related to Ca^2+^ homeostasis, VSMCs profile, Ca^2+^-mediated mineralization and VSMCs-ECM crosstalk via integrins signaling (**Fig. 5-6**). Physiologically, NCKX4-mediated Ca^2+^ clearance ensures that cytosolic Ca^2+^ levels are tightly regulated, preventing Ca^2+^ overload and sustaining the activation of Ca^2+^-dependent signaling cascades ^[12, 89]^. Ca^2+^ handling and calcification are critical in the pathogenesis of vascular calcification, plasticity and aging ^[90–92]^. Our pathway analysis identified Ca^2+^-mediated mineralization pathways with different expression levels particularly in the aorta of young *Nckx4^−/−^*mice. Several differentially expressed genes were directly linked to mineralization-induced osteogenic differentiation (*Bmp2* and *Bmper*), activation of BMP/TGF-β (*Smad6* and *Smad7*) and Wnt/Ca^2+^ (*Wnt5a/b* and *Wnt9a*), and osteogenic reprogramming factors (*Sox5/6* and *Runx1*) (**Fig. 5-6**). These genes are linked to VSMCs mineralization both in murine models and human vascular diseases ^[43, 86, 92]^. Pathways associated with BMP signaling are elevated in calcified human valves and arteries ^[93, 94]^, within Wnt-induced calcifications of VSMCs involving Wnt5a upregulation and cooperation of BMP/Runx programs to promote osteogenic factors ^[95, 96]^. By contrast, loss of inhibitory Smads (*Smad6/7*), oxidative stress markers and changes in senescence (*Cdkn1a*, Gpx and *Sod*) genes further predispose VSMCs to mineralization in vessels ^[37, 43, 97, 98]^. This links senescence-associated changes to calcification propensity. Together, these studies corroborate our observation that NCKX4 deficiency and aging converge on canonical osteogenic pathways, providing a potential route from impaired Ca²⁺ clearance to early vascular calcification.

An impaired capacity to remove cytosolic Ca^2+^ via genetic deletion of NCKX4 in young mice, or by natural age-related reduction of NCKX4 expression could result in prolonged and sustained high [Ca^2+^]_cyt_ levels. This may initiate maladaptive inside-out integrin signaling, altering VSMCs contractility and adhesion dynamics, calcification processes and ECM turnover. In fact, transient increase in cytosolic Ca^2+^ levels ranging from 400 nM to 1 μM is a potent regulator of integrin inside-out activation and cellular dynamics ^[8, 38, 99–102]^. Studies in primary murine VSMCs, human leukocytes and HL-60 cell line demonstrated that increases in [Ca^2+^]_cyt_ levels is necessary for inside-out activation of β integrins and ECM remodeling ^[8, 38, 101, 103]^. Additionally, sustained Ca²⁺ elevations also activate calpain proteases that cleave talin/FAK/paxillin components, altering adhesion stability and turnover ^[99, 103, 104]^. These Ca²⁺-dependent processes could chronically produce maladaptive ECM remodeling, VSMCs phenotypic switching, calcification and loss of proper integrin anchorage. These potential targets and mechanisms are supported by our RNA-seq, where NCKX4 loss precipitates a premature switch in the integrin-ECM interaction associated with Ca^2+^ signaling and mineralization pathways (**Fig. 5-6**). We showed that protective anchorage integrins (*Itga8*, *Itgav*, and *Itga4*) and microfibril-stabilizing proteins (Adamtsl1/10/17) and balanced proteolysis (*Timp2/3*) were markedly downregulated, while pro-fibrotic integrins (*Col8a1*, *Itga5/11* and *Itgb8*) were upregulated in aorta from young *Nckx4^−/−^* mice.

A critical aspect of vascular homeostasis is the balance between firm integrin anchorage and controlled ECM proteolysis, which allows remodeling without compromising wall stability ^[7, 102]^. In vascular wall integrins such as α5β1, αvβ3, αvβ5, and α8β1 anchor VSMCs to fibronectin and other ECM components, ensuring proper adhesion, mechanotransduction, and survival signaling ^[7, 8]^. Integrin-mediated adhesion complexes recruit focal adhesion proteins and cross-talk with matrix metalloproteinases (MMPs) and tissue inhibitors of metalloproteinases (TIMPs), ensuring that ECM degradation does not exceed synthesis ^[7, 36, 105]^. Mice lacking α8 integrin (*Itga8^-/-^*) show vascular lesions, medial thickening, and increased plaque formation ^[106]^. A recent report from Martínez et al. demonstrated that VSMC-specific knockout of *Itga8* causes early aortic aneurysms ^[107]^, indicating that *Itga8*-expressing VSMCs are critical for maintaining ECM structure and preventing remodeling. In addition, loss of integrin α5 was observed in vessels from aged (∼22-24-month-old) C57Bl/6 mice, suggesting a lower firm integrin anchorage ^[108]^. The deletion of *Timp3* results in increased proteolytic activity of MMPs with degraded elastin, disrupted ECM, decreased collagen/elastin protein, and wall structure breakdown in adult C57Bl/6 mice ^[109]^. This shows that lack of protease inhibition causes ECM degradation and remodeling. Our results showed that the loss of NCKX4 in young mice significantly disrupted the balance between firm integrin anchorage and controlled ECM proteolysis, suggesting that these pathways are potentially involved in the early aortic remodeling. This resembles aged WT profile and implies that impaired Ca²⁺ clearance disrupts the balance between firm integrin anchorage and controlled proteolysis, being critical for vascular integrity. Such imbalance promotes ECM disorganization, elastic fiber fragmentation, and maladaptive stiffening, thereby linking Ca²⁺-dependent signaling to early vascular aging.

## 5. CONCLUSION

Our study identifies NCKX4 as a key regulator of Ca^2+^ clearance in VSMCs and aortic vascular integrity during aging. We demonstrate that genetic deletion or age-related reduction of NCKX4 accelerates aortic remodeling, characterized by impaired Ca^2+^ removal, elastic fiber fragmentation, and excessive collagen deposition, all hallmarks of vascular aging. NCKX4 loss triggers premature activation of Ca^2+^-dependent pathways that converge on integrin signaling, ECM remodeling, and mineralization processes. Transcriptomic profiling revealed early imbalance between firm integrin anchorage and controlled proteolysis provides a mechanistic link between impaired Ca^2+^ clearance and maladaptive vascular remodeling. Therefore, our findings position NCKX4 as a new Ca^2+^ homeostasis regulating protein involved in vascular aging, whereby its decline promotes premature stiffening and calcification through Ca^2+^-integrin-mediated pathways. Targeting NCKX4 or its downstream signaling may represent a novel therapeutic avenue to early detect, preserve vascular integrity and mitigate age-related risk of CVDs development.

## AUTHOR CONTRIBUTIONS

**Guilherme H. Souza Bomfim =** Conceptualization, formal analysis, investigation, experimentation, methodology, validation, writing of the draft, review and editing; **Kesava Asam =** Formal analysis, investigation, experimentation and validation; **Nish Patel =** Formal analysis, experimentation and validation; **Kristen Rosenberg =** Formal analysis, investigation, experimentation and validation; **Erna Mitaishvili =** Formal analysis, experimentation and validation; **Talita Aguiar =** Formal analysis, experimentation and validation**; Emmanuel Zorn =** Funding acquisition, formal analysis and investigation; **Bradley Aouizerat =** Formal analysis, funding acquisition, methodology, visualization; **Ravichandran Ramasamy =** Conceptualization, formal analysis and visualization; **Rodrigo S. Lacruz =** Conceptualization, supervision, funding acquisition, methodology, validation, writing of the draft, review and editing.

## ACKNOWLEDGMENTS

This research was supported by the Department of Molecular Pathobiology Accelerator Award (P01) 2025, New York University College of Dentistry. Rodrigo S. Lacruz was supported by intramural funding from New York University (NYU, grant no. MA003). We thank Dr. Haiqing Zhao (John Hopkins) for generously providing NCKX4 knock-out (*Nckx4^-/-^*) mice.

We thank NYU Langone’s Genome Technology Center (RRID: SCR_017929) for instruments and services of RNA sequencing (RNA-seq). We thank NYU Langone’s Experimental Pathology Research Laboratory (RRID:SCR_017928) for histological proceeds. This laboratory is partially supported by the Cancer Center Support Grant P30CA016087 at NYU Langone.

## DATA AVAILABILITY STATEMENT

The data that support the findings of this study are available in the methods and/or supplementary material of this article.

## REFERENCES

1. Tsao, C.W., et al., Heart Disease and Stroke Statistics-2022 Update: A Report From the American Heart Association. Circulation, 2022. 145(8): p. e153–e639.

2. Goetzman, E., et al., Complex II Biology in Aging, Health, and Disease. Antioxidants (Basel), 2023. 12(7).

3. Rodgers, J.L., et al., Cardiovascular Risks Associated with Gender and Aging. J Cardiovasc Dev Dis, 2019. 6(2).

4. Garcia, M., et al., Cardiovascular Disease in Women: Clinical Perspectives. Circ Res, 2016. 118(8): p. 1273–93.

5. Harraz, O.F. and L.J. Jensen, Aging, calcium channel signaling and vascular tone. Mech Ageing Dev, 2020. 191: p. 111336.

6. Harraz, O.F. and L.J. Jensen, Vascular calcium signalling and ageing. J Physiol, 2021. 599(24): p. 5361–5377.

7. Jain, M. and A.K. Chauhan, Role of Integrins in Modulating Smooth Muscle Cell Plasticity and Vascular Remodeling: From Expression to Therapeutic Implications. Cells, 2022. 11(4).

8. Zhu, Y., et al., Calcium in Vascular Smooth Muscle Cell Elasticity and Adhesion: Novel Insights Into the Mechanism of Action. Front Physiol, 2019. 10: p. 852.

9. Guo, J., et al., Aging and aging-related diseases: from molecular mechanisms to interventions and treatments. Signal Transduct Target Ther, 2022. 7(1): p. 391.

10. Climie, R.E., et al., Vascular ageing: moving from bench towards bedside. Eur J Prev Cardiol, 2023. 30(11): p. 1101–1117.

11. Felbel, J., et al., Regulation of cytosolic calcium by cAMP and cGMP in freshly isolated smooth muscle cells from bovine trachea. J Biol Chem, 1988. 263(32): p. 16764–71.

12. Berridge, M.J., M.D. Bootman, and H.L. Roderick, Calcium signalling: dynamics, homeostasis and remodelling. Nat Rev Mol Cell Biol, 2003. 4(7): p. 517–29.

13. Reid, I.R., S.M. Birstow, and M.J. Bolland, Calcium and Cardiovascular Disease. Endocrinol Metab (Seoul), 2017. 32(3): p. 339–349.

14. Wagenseil, J.E. and R.P. Mecham, Vascular extracellular matrix and arterial mechanics. Physiol Rev, 2009. 89(3): p. 957–89.

15. Owens, G.K., M.S. Kumar, and B.R. Wamhoff, Molecular regulation of vascular smooth muscle cell differentiation in development and disease. Physiol Rev, 2004. 84(3): p. 767–801.

16. Majesky, M.W., Developmental basis of vascular smooth muscle diversity. Arterioscler Thromb Vasc Biol, 2007. 27(6): p. 1248–58.

17. Altimimi, H.F. and P.P. Schnetkamp, Na+/Ca2+-K+ exchangers (NCKX): functional properties and physiological roles. Channels (Austin), 2007. 1(2): p. 62–9.

18. Jalloul, A.H., et al., Structure-function relationships of K(+)-dependent Na(+)/Ca(2+) exchangers (NCKX). Cell Calcium, 2020. 86: p. 102153.

19. Li, X.F., A.S. Kraev, and J. Lytton, Molecular cloning of a fourth member of the potassium-dependent sodium-calcium exchanger gene family, NCKX4. J Biol Chem, 2002. 277(50): p. 48410–7.

20. Zhang, S., et al., Role of Na+/Ca2+ exchange in regulating cytosolic Ca2+ in cultured human pulmonary artery smooth muscle cells. Am J Physiol Cell Physiol, 2005. 288(2): p. C245–52.

21. Al-Khannaq, M. and J. Lytton, Regulation of K(+)-Dependent Na(+)/Ca(2+)-Exchangers (NCKX). Int J Mol Sci, 2022. 24(1).

22. Hassan, M.T. and J. Lytton, Potassium-dependent sodium-calcium exchanger (NCKX) isoforms and neuronal function. Cell Calcium, 2020. 86: p. 102135.

23. Dong, H., et al., Novel role for K+-dependent Na+/Ca2+ exchangers in regulation of cytoplasmic free Ca2+ and contractility in arterial smooth muscle. Am J Physiol Heart Circ Physiol, 2006. 291(3): p. H1226–35.

24. Reeves, J.P. and M. Condrescu, Lanthanum is transported by the sodium/calcium exchanger and regulates its activity. Am J Physiol Cell Physiol, 2003. 285(4): p. C763–70.

25. Lenart, B., et al., Na-K-Cl cotransporter-mediated intracellular Na+ accumulation affects Ca2+ signaling in astrocytes in an in vitro ischemic model. J Neurosci, 2004. 24(43): p. 9585–97.

26. Palty, R., et al., Lithium-calcium exchange is mediated by a distinct potassium-independent sodium-calcium exchanger. J Biol Chem, 2004. 279(24): p. 25234–40.

27. Hilge, M., Ca2+ regulation of ion transport in the Na+/Ca2+ exchanger. J Biol Chem, 2012. 287(38): p. 31641–9.

28. Annunziato, L., et al., New perspectives for selective NCX activators in neurodegenerative diseases. Cell Calcium, 2020. 87: p. 102170.

29. Verkhratsky, A., et al., Crosslink between calcium and sodium signalling. Exp Physiol, 2018. 103(2): p. 157–169.

30. Berra-Romani, R., et al., The mechanism of injury-induced intracellular calcium concentration oscillations in the endothelium of excised rat aorta. J Vasc Res, 2012. 49(1): p. 65–76.

31. Khananshvili, D., Basic and editing mechanisms underlying ion transport and regulation in NCX variants. Cell Calcium, 2020. 85: p. 102131.

32. Wang, Y., J. Shi, and X. Tong, Cross-Talk between Mechanosensitive Ion Channels and Calcium Regulatory Proteins in Cardiovascular Health and Disease. Int J Mol Sci, 2021. 22(16).

33. Lacolley, P., V. Regnault, and A.P. Avolio, Smooth muscle cell and arterial aging: basic and clinical aspects. Cardiovasc Res, 2018. 114(4): p. 513–528.

34. Ojha, K.R., et al., Age-Associated Dysregulation of Integrin Function in Vascular Smooth Muscle. Front Physiol, 2022. 13: p. 913673.

35. Sjaastad, M.D. and W.J. Nelson, Integrin-mediated calcium signaling and regulation of cell adhesion by intracellular calcium. Bioessays, 1997. 19(1): p. 47–55.

36. Schwartz, M.A., Integrins and extracellular matrix in mechanotransduction. Cold Spring Harb Perspect Biol, 2010. 2(12): p. a005066.

37. Cao, G., et al., How vascular smooth muscle cell phenotype switching contributes to vascular disease. Cell Commun Signal, 2022. 20(1): p. 180.

38. Huang, H., et al., A Calcium Mediated Mechanism Coordinating Vascular Smooth Muscle Cell Adhesion During KCl Activation. Front Physiol, 2018. 9: p. 1810.

39. Zhu, S., et al., Cell signaling and transcriptional regulation of osteoblast lineage commitment, differentiation, bone formation, and homeostasis. Cell Discov, 2024. 10(1): p. 71.

40. Adeyemo, A., et al., A genome-wide association study of hypertension and blood pressure in African Americans. PLoS Genet, 2009. 5(7): p. e1000564.

41. Kidambi, S., et al., Non-replication study of a genome-wide association study for hypertension and blood pressure in African Americans. BMC Med Genet, 2012. 13: p. 27.

42. Gui, T., et al., MicroRNAs that target Ca(2+) transporters are involved in vascular smooth muscle cell calcification. Lab Invest, 2012. 92(9): p. 1250–9.

43. Sutton, N.R., et al., Molecular Mechanisms of Vascular Health: Insights From Vascular Aging and Calcification. Arterioscler Thromb Vasc Biol, 2023. 43(1): p. 15–29.

44. McHugh, D. and J. Gil, Senescence and aging: Causes, consequences, and therapeutic avenues. J Cell Biol, 2018. 217(1): p. 65–77.

45. Folgueras, A.R., et al., Mouse Models to Disentangle the Hallmarks of Human Aging. Circ Res, 2018. 123(7): p. 905–924.

46. Mansfield, L., et al., Emerging insights in senescence: pathways from preclinical models to therapeutic innovations. NPJ Aging, 2024. 10(1): p. 53.

47. De Moudt, S., et al., Progressive aortic stiffness in aging C57Bl/6 mice displays altered contractile behaviour and extracellular matrix changes. Commun Biol, 2022. 5(1): p. 605.

48. Yanai, S. and S. Endo, Functional Aging in Male C57BL/6J Mice Across the Life-Span: A Systematic Behavioral Analysis of Motor, Emotional, and Memory Function to Define an Aging Phenotype. Front Aging Neurosci, 2021. 13: p. 697621.

49. Li, L., et al., Klotho Ameliorates Vascular Calcification via Promoting Autophagy. Oxid Med Cell Longev, 2022. 2022: p. 7192507.

50. Stephan, A.B., et al., The Na(+)/Ca(2+) exchanger NCKX4 governs termination and adaptation of the mammalian olfactory response. Nat Neurosci, 2011. 15(1): p. 131–7.

51. Adhikari, N., et al., Guidelines for the isolation and characterization of murine vascular smooth muscle cells. A report from the International Society of Cardiovascular Translational Research. J Cardiovasc Transl Res, 2015. 8(3): p. 158–63.

52. Méndez-López, I., et al., Altered mitochondrial function, capacitative calcium entry and contractions in the aorta of hypertensive rats. J Hypertens, 2017. 35(8): p. 1594–1608.

53. Souza Bomfim, G.H., et al., Na+/Ca2+ exchange in enamel cells is dominated by the K+-dependent NCKX exchanger. J Gen Physiol, 2024. 156(1).

54. Aguiar, T., et al., Development of high-titer class-switched antibody responses to phosphorylated amino acids is prevalent in pancreatic ductal adenocarcinoma. Front Immunol, 2025. 16: p. 1501943.

55. Chevalier, C., et al., Primary mouse osteoblast and osteoclast culturing and analysis. STAR Protoc, 2021. 2(2): p. 100452.

56. Souza Bomfim, G.H., et al., Functional Upregulation of STIM-1/Orai-1-Mediated Store-Operated Ca2+ Contributing to the Hypertension Development Elicited by Chronic EtOH Consumption. Curr Vasc Pharmacol, 2017. 15(3): p. 265–281.

57. Son, G.Y., et al., The Ca(2+) channel ORAI1 is a regulator of oral cancer growth and nociceptive pain. Sci Signal, 2023. 16(801): p. eadf9535.

58. Aulestia, F.J., et al., Fluoride exposure alters Ca(2+) signaling and mitochondrial function in enamel cells. Sci Signal, 2020. 13(619).

59. Bomfim, G.H.S., M. Giacomello, and R.S. Lacruz, PMCA Ca(2+) clearance in dental enamel cells depends on the magnitude of cytosolic Ca(2). Faseb j, 2023. 37(1): p. e22679.

60. Souza Bomfim, G.H., et al., TRPM7 activation potentiates SOCE in enamel cells but requires ORAI. Cell Calcium, 2020. 87: p. 102187.

61. Lytton, J., et al., K+-dependent Na+/Ca2+ exchangers in the brain. Ann N Y Acad Sci, 2002. 976: p. 382–93.

62. Zha, Y., et al., Senescence in Vascular Smooth Muscle Cells and Atherosclerosis. Front Cardiovasc Med, 2022. 9: p. 910580.

63. Oloizia, B. and R.J. Paul, Ca2+ clearance and contractility in vascular smooth muscle: evidence from gene-altered murine models. J Mol Cell Cardiol, 2008. 45(3): p. 347–62.

64. Liu, Z. and R.A. Khalil, Evolving mechanisms of vascular smooth muscle contraction highlight key targets in vascular disease. Biochem Pharmacol, 2018. 153: p. 91–122.

65. Lin, Y. and Z. Sun, Klotho deficiency-induced arterial calcification involves osteoblastic transition of VSMCs and activation of BMP signaling. J Cell Physiol, 2022. 237(1): p. 720–729.

66. Park, J.H., et al., Relationship between arterial calcification and bone loss in a new combined model rat by ovariectomy and vitamin D(3) plus nicotine. Calcif Tissue Int, 2008. 83(3): p. 192–201.

67. Hammad, S.K., et al., Resveratrol Ameliorates Aortic Calcification in Ovariectomized Rats via SIRT1 Signaling. Curr Issues Mol Biol, 2021. 43(2): p. 1057–1071.

68. Kilanowski-Doroh, I.M., et al., Ovariectomy-Induced Arterial Stiffening Differs From Vascular Aging and Is Reversed by GPER Activation. Hypertension, 2024. 81(5): p. e51–e62.

69. Cecelja, M., et al., Arterial stiffening relates to arterial calcification but not to noncalcified atheroma in women. A twin study. J Am Coll Cardiol, 2011. 57(13): p. 1480–6.

70. Vallée, A., Menopause and arterial stiffness index: insights from the women’s UK Biobank cohort. Maturitas, 2025. 198: p. 108608.

71. Samargandy, S., et al., Arterial Stiffness Accelerates Within 1 Year of the Final Menstrual Period: The SWAN Heart Study. Arterioscler Thromb Vasc Biol, 2020. 40(4): p. 1001–1008.

72. Berridge, M.J., Calcium microdomains: organization and function. Cell Calcium, 2006. 40(5-6): p. 405–12.

73. Bronner, F., Extracellular and intracellular regulation of calcium homeostasis. ScientificWorldJournal, 2001. 1: p. 919–25.

74. Kapustin, A.N., et al., Calcium regulates key components of vascular smooth muscle cell-derived matrix vesicles to enhance mineralization. Circ Res, 2011. 109(1): p. e1–12.

75. Lacolley, P., et al., Vascular Smooth Muscle Cells and Arterial Stiffening: Relevance in Development, Aging, and Disease. Physiol Rev, 2017. 97(4): p. 1555–1617.

76. Mammoto, A., K. Matus, and T. Mammoto, Extracellular Matrix in Aging Aorta. Front Cell Dev Biol, 2022. 10: p. 822561.

77. Vanalderwiert, L., et al., Exploring aortic stiffness in aging mice: a comprehensive methodological overview. Aging (Albany NY), 2024. 17(2): p. 280–307.

78. Jegger, D., et al., Effects of an aging vascular model on healthy and diseased hearts. Am J Physiol Heart Circ Physiol, 2007. 293(3): p. H1334–43.

79. Hawes, J.Z., et al., Elastin haploinsufficiency in mice has divergent effects on arterial remodeling with aging depending on sex. Am J Physiol Heart Circ Physiol, 2020. 319(6): p. H1398–h1408.

80. Intengan, H.D. and E.L. Schiffrin, Vascular remodeling in hypertension: roles of apoptosis, inflammation, and fibrosis. Hypertension, 2001. 38(3 Pt 2): p. 581–7.

81. Kim, S.H., et al., Age-associated proinflammatory elastic fiber remodeling in large arteries. Mech Ageing Dev, 2021. 196: p. 111490.

82. Xu, X., et al., Age-related Impairment of Vascular Structure and Functions. Aging Dis, 2017. 8(5): p. 590–610.

83. Christensen, K.B., et al., MFAP4-Deficiency Aggravates Age-Induced Changes in Resistance Artery Structure, While Ameliorating Hypertension. Hypertension, 2024. 81(6): p. 1308–1319.

84. Cattell, M.A., J.C. Anderson, and P.S. Hasleton, Age-related changes in amounts and concentrations of collagen and elastin in normotensive human thoracic aorta. Clin Chim Acta, 1996. 245(1): p. 73–84.

85. Shahbad, R., et al., Effects of age, elastin density, and glycosaminoglycan accumulation on the delamination strength of human thoracic and abdominal aortas. Acta Biomater, 2024. 189: p. 413–426.

86. Cecelja, M., et al., Multimodality imaging of subclinical aortic atherosclerosis: relation of aortic stiffness to calcification and plaque in female twins. Hypertension, 2013. 61(3): p. 609–14.

87. You, A.Y.F., et al., Raman spectroscopy imaging reveals interplay between atherosclerosis and medial calcification in the human aorta. Sci Adv, 2017. 3(12): p. e1701156.

88. Schlatmann, T.J. and A.E. Becker, Histologic changes in the normal aging aorta: implications for dissecting aortic aneurysm. Am J Cardiol, 1977. 39(1): p. 13–20.

89. Bootman, M.D., Calcium signaling. Cold Spring Harb Perspect Biol, 2012. 4(7): p. a011171.

90. Jono, S., et al., Phosphate regulation of vascular smooth muscle cell calcification. Circ Res, 2000. 87(7): p. E10–7.

91. Petsophonsakul, P., et al., Role of Vascular Smooth Muscle Cell Phenotypic Switching and Calcification in Aortic Aneurysm Formation. Arterioscler Thromb Vasc Biol, 2019. 39(7): p. 1351–1368.

92. Proudfoot, D., Calcium Signaling and Tissue Calcification. Cold Spring Harb Perspect Biol, 2019. 11(10).

93. Gomez-Stallons, M.V., et al., Bone Morphogenetic Protein Signaling Is Required for Aortic Valve Calcification. Arterioscler Thromb Vasc Biol, 2016. 36(7): p. 1398–405.

94. Li, X., H.Y. Yang, and C.M. Giachelli, BMP-2 promotes phosphate uptake, phenotypic modulation, and calcification of human vascular smooth muscle cells. Atherosclerosis, 2008. 199(2): p. 271–7.

95. Freise, C., N. Kretzschmar, and U. Querfeld, Wnt signaling contributes to vascular calcification by induction of matrix metalloproteinases. BMC Cardiovasc Disord, 2016. 16(1): p. 185.

96. Bundy, K., J. Boone, and C.L. Simpson, Wnt Signaling in Vascular Calcification. Front Cardiovasc Med, 2021. 8: p. 708470.

97. Leopold, J.A., *Vascular calcification: an age-old problem of old age*, in *Circulation*. 2013: United States. p. 2380–2.

98. Li, X., et al., Protective Role of Smad6 in Inflammation-Induced Valvular Cell Calcification. J Cell Biochem, 2015. 116(10): p. 2354–64.

99. Dewitt, S. and M.B. Hallett, Cytosolic free Ca(2+) changes and calpain activation are required for beta integrin-accelerated phagocytosis by human neutrophils. J Cell Biol, 2002. 159(1): p. 181–9.

100. Dixit, N., et al., Migrational guidance of neutrophils is mechanotransduced via high-affinity LFA-1 and calcium flux. J Immunol, 2011. 187(1): p. 472–81.

101. Schaff, U.Y., et al., Calcium flux in neutrophils synchronizes beta2 integrin adhesive and signaling events that guide inflammatory recruitment. Ann Biomed Eng, 2008. 36(4): p. 632–46.

102. Sun, Z., et al., Extracellular matrix-specific focal adhesions in vascular smooth muscle produce mechanically active adhesion sites. Am J Physiol Cell Physiol, 2008. 295(1): p. C268–78.

103. Kerstein, P.C., K.M. Patel, and T.M. Gomez, Calpain-Mediated Proteolysis of Talin and FAK Regulates Adhesion Dynamics Necessary for Axon Guidance. J Neurosci, 2017. 37(6): p. 1568–1580.

104. Saxena, M., et al., Force-Induced Calpain Cleavage of Talin Is Critical for Growth, Adhesion Development, and Rigidity Sensing. Nano Lett, 2017. 17(12): p. 7242–7251.

105. Pang, X., et al., Targeting integrin pathways: mechanisms and advances in therapy. Signal Transduct Target Ther, 2023. 8(1): p. 1.

106. Menendez-Castro, C., et al., *Under-expression of* α*8 integrin aggravates experimental atherosclerosis*. J Pathol, 2015. 236(1): p. 5–16.

107. Arévalo Martínez, M., et al., Vascular smooth muscle-specific YAP/TAZ deletion triggers aneurysm development in mouse aorta. JCI Insight, 2023. 8(17).

108. Wei, J., et al., Aging Impairs Renal Autoregulation in Mice. Hypertension, 2020. 75(2): p. 405–412.

109. Basu, R., et al., Loss of Timp3 gene leads to abdominal aortic aneurysm formation in response to angiotensin II. J Biol Chem, 2012. 287(53): p. 44083–96.

