## Supplementary Materials for "Age-Related Decline in NCKX4-Mediated Calcium Clearance Accelerates Aortic Remodeling and Drives Early Vascular Aging"

Souza Bomfim, GH. ORCID: <https://orcid.org/0000-0001-9454-7266>

Lacruz, RS. ORCID: <https://orcid.org/0000-0002-0776-6143>

**Includes:**

**Figure S1.** Genotyping of mice to detect the wild-type (WT) allele and homozygous *Nckx4* allele deletion (*Nckx4^−/−^*).

**Figure S2.** Gene expression of *Nckx4* in brain, heart, stomach and liver isolated from young (Y, 12-15 weeks) and aged (A, 72-78 weeks) male and female wild-type (WT) mice.

**Figure S3.** Uncropped and unprocessed immunoblot membranes of NCKX4 and GAPDH expression in thoracic aorta isolated from young (Y, 12-15 weeks) and aged (A, 72-78 weeks) male and female wild-type (WT) mice.

**Figure S4.** Immunofluorescence of NCKX4 in aortic tissue and primary VSMCs from female mouse lacking NCKX4 (*Nckx4^-/-^*).

**Figure S5.** Validation of NCKX4 and specificity of the secondary antibody in aortic tissue and primary VSMCs from wild-type (WT) female.

**Figure S6.** Chemical and pharmacological validation of the NCKX/NCX activity in Ca^2+^ influx reverse mode in primary VSMCs from young (Y, 12-15 weeks) male and female wild-type (WT) mice loaded with the Ca²⁺ probe Fura-2-AM.

**Figure S7.** Alizarin Red-S staining in primary VSMCs from young (Y, 12-15 weeks) and aged (A, 72-78 weeks) male and female *Nckx4* knockout (*Nckx4^-/-^*) and wild-type (WT) mice. The VSMCs were culture in a non-procalcifying control media (high glucose regular DMEM media) for 16 days.

**Figure S8.** Principal Component Analysis (PCA) of bulk RNA-seq data of aortic tissue from young (Y, 12-15 weeks) and aged (A, 72-78 weeks) female wild-type (WT) and *Nckx4^-/-^* mice.

**Figure S9.** Volcano plot of differential gene expression of bulk RNA-seq data of aortic tissue from young (Y, 12-15 weeks) and aged (A, 72-78 weeks) female wild-type (WT) and *Nckx4^-/-^* mice.

**Figure S10.** Differential expression analysis of pathways identified genes with significantly different expression using gene regulatory network representation.

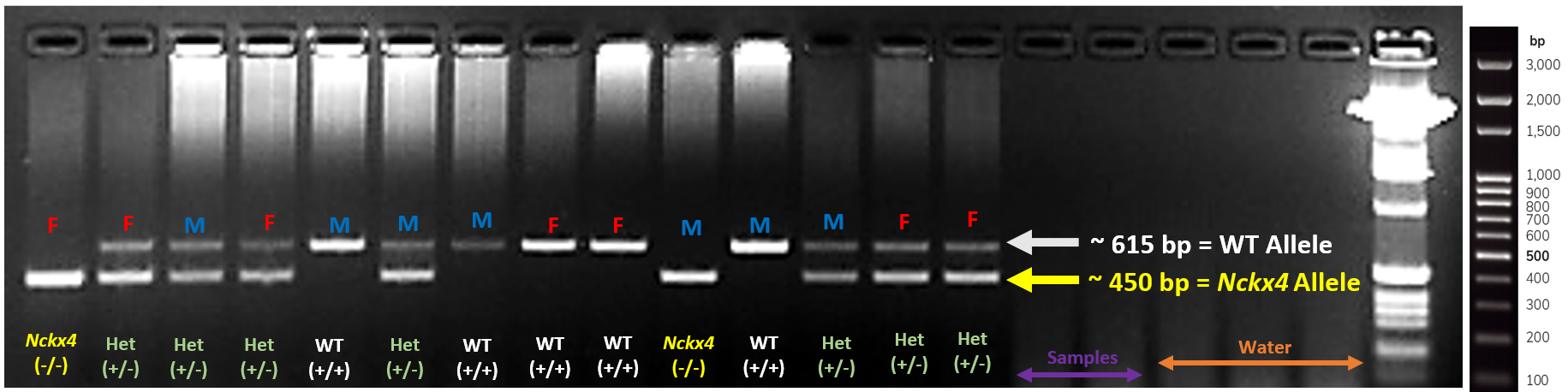

**Figure S1. Genotyping of mice to detect the wild-type (WT) allele and homozygous *Nckx4* allele deletion (*Nckx4^−/−^*)**. Representative genotyping obtained from *Nckx4* heterozygous (*+/-*) offspring breeding using mouse tail DNA and PCR primers specific for the WT allele and the recombined *Nckx4* allele in order to validate the animals used in this study. Primers AAAGGAAATGAAGAGAAGGC (AS67) and TCATCTCATAAGAAGCCCAG (AS68), were used to genotype the *Nckx4* allele, with expected band sizes being 615 bp for wild-type (WT) and 450 bp for the knocked-out (*Nckx4^-/-^*) exon. PCR conditions: Initial denaturation at 95°C for 10 min, followed by 30 cycles of 95°C for 30 s, annealing at 55°C for 30 s, and elongation at 72°C for 60 s, with a final elongation step at 72°C for 10 min. Mice heterozygous for the *Nckx4* allele deletion (*Nckx4^+/−^*) were used as a control to confirm PCR primer specificity.

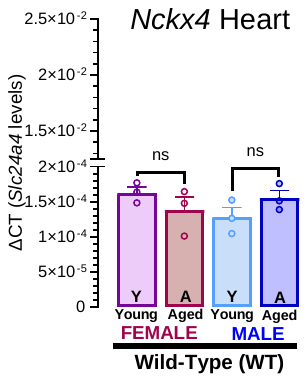

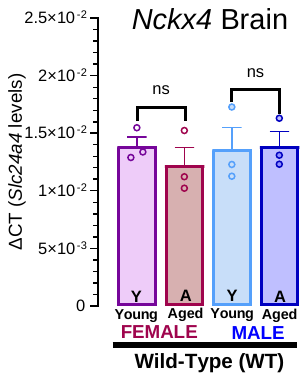

**B**

**A**

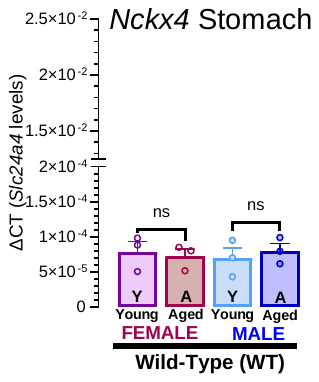

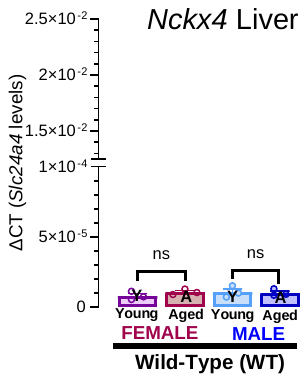

**D**

**C**

**Figure S2.** Gene expression of *Nckx4* in brain (**A**), heart (**B**), stomach (**C**) and liver (**D**) isolated from young (Y, 12-15 weeks) and aged (A, 72-78 weeks) male and female wild-type (WT) mice. mRNA levels of *Slc24a4* (*Nckx4*) were analyzed by qRT-PCR using the 2^−ΔCT^ method using β-actin as the housekeeping control. The results were analyzed by one-way ANOVA followed by Tukey’s multiple comparison post-hoc test. Significance was accepted at n.s, non-significant.

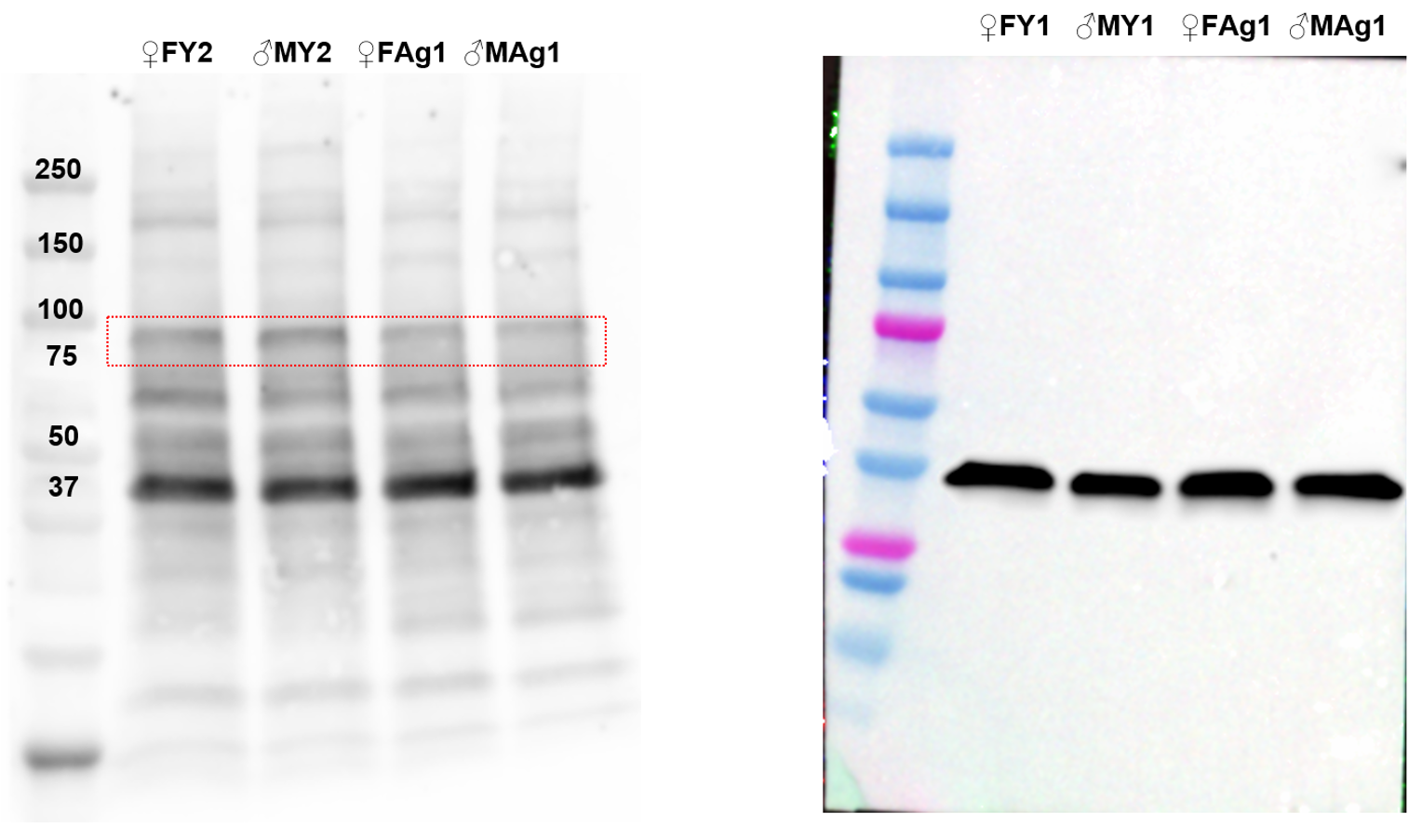

**A**

**B**

**Figure S3.** Uncropped and unprocessed immunoblot membranes of *Slc24a4* (*Nckx4*) (**A**) and GAPDH (**B**) expression in each sample, as a control, in thoracic aorta isolated from young (Y, 12-15 weeks) and aged (A, 72-78 weeks) male and female wild-type (WT) mice. FY = female young samples; MY = male young samples; FAg = female aged samples; MAg = male aged samples. *Slc24a4* (*Nckx4*) = ~65-75 kDa. GAPDH = ~38 kDa.

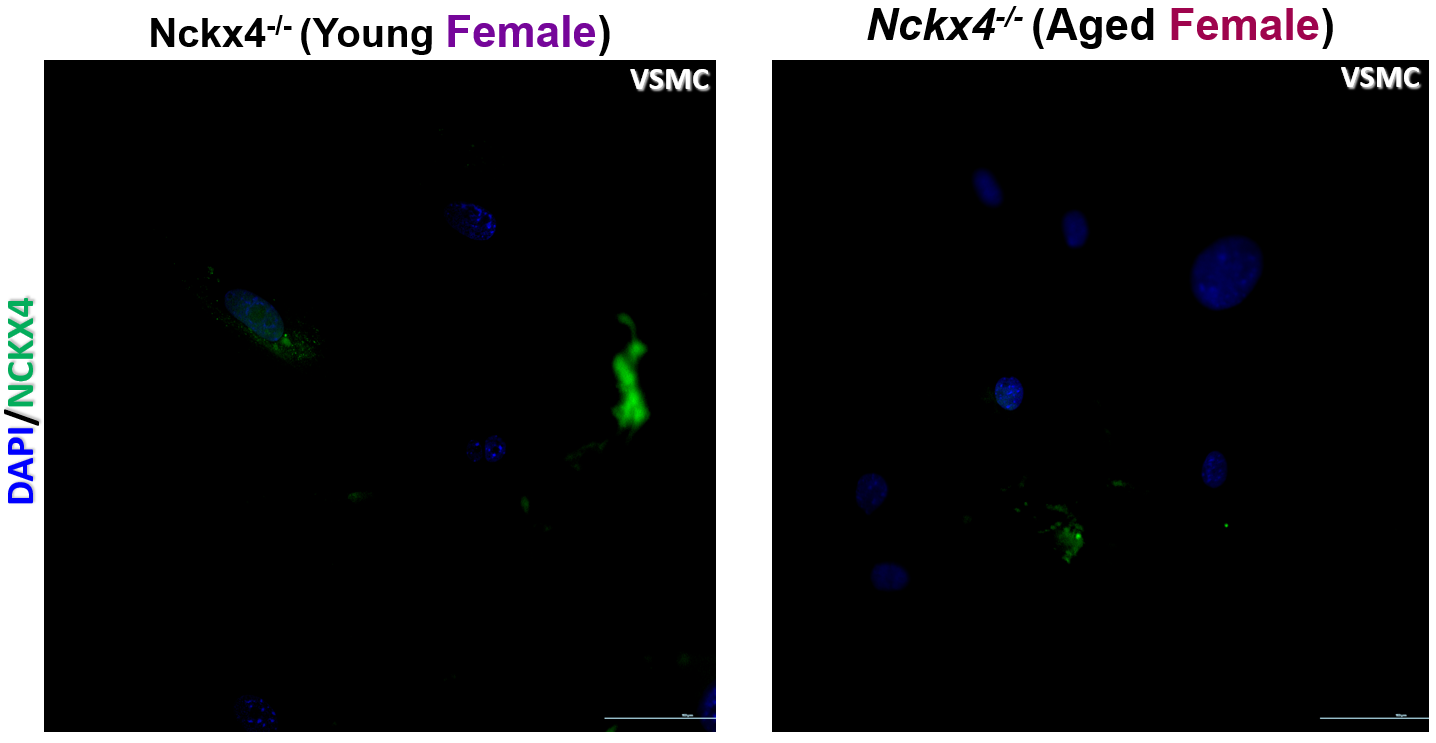

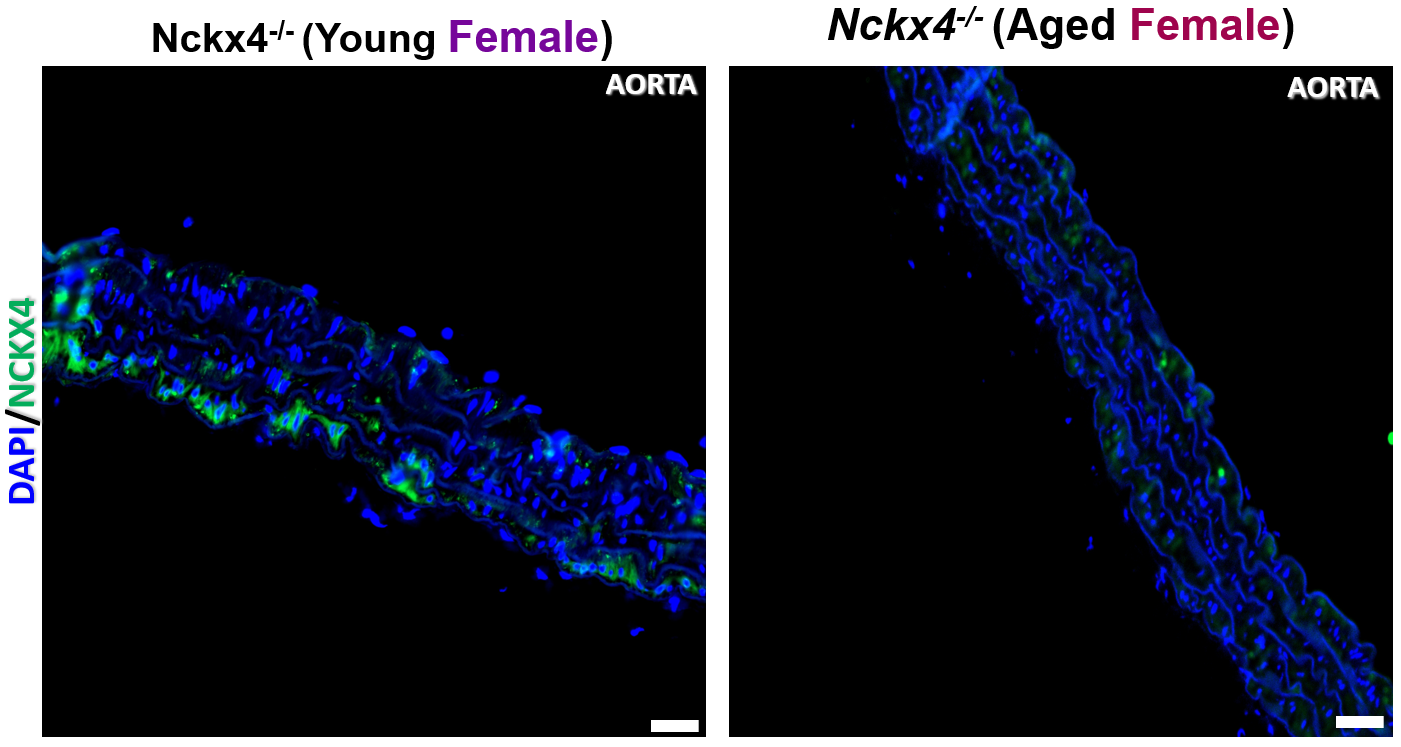

**A**

**B**

**C**

**D**

**Figure S4.** Immunofluorescence of NCKX4 in aortic tissue and primary VSMCs from female mouse lacking NCKX4 (*Nckx4^-/-^*). Immunofluorescence staining of aorta (**A-B**) and VSMCs (**C-D**) from young (Y, 12-15 weeks) and aged (A, 72-78 weeks) female *Nckx4^-/-^*) mice. Anti-NCKX4 is showed in green, autofluorescent of elastin and elastic fibers at blue excitation light (~340-400 nm), and nuclear staining of VSMCs with DAPI/Hoechst showed in blue.

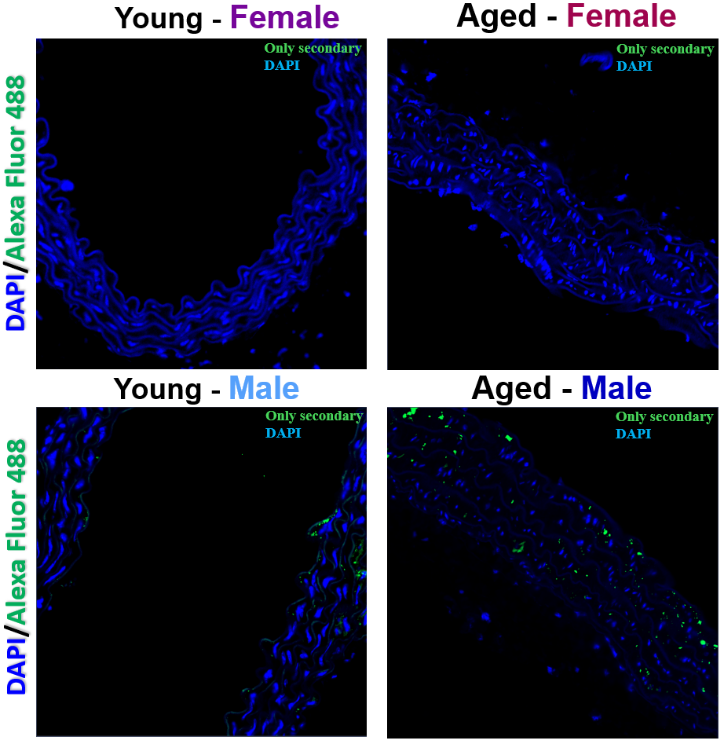

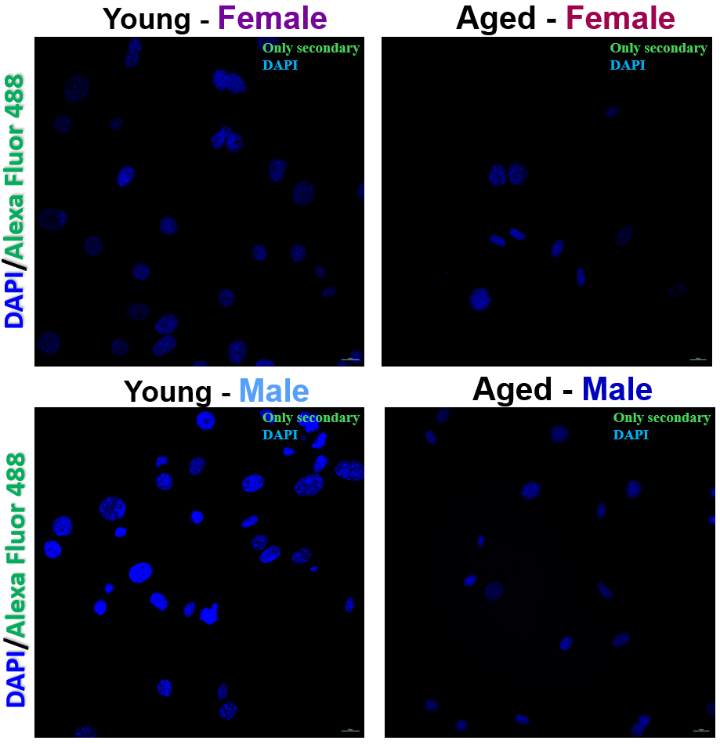

**A**

**B**

**Figure S5.** Validation of NCKX4 and specificity of the secondary antibody. Immunofluorescence of NCKX4 in aortic tissue (A) and primary VSMCs (B) from young (Y, 12-15 weeks) and aged (A, 72-78 weeks) female wild-type (WT) mice. Only Goat anti-mouse Alexa Fluor 488 1:800 (green), autofluorescent of elastin and elastic fibers at blue excitation light (~340-400 nm), and nuclear staining of VSMCs with DAPI/Hoechst showed in blue.

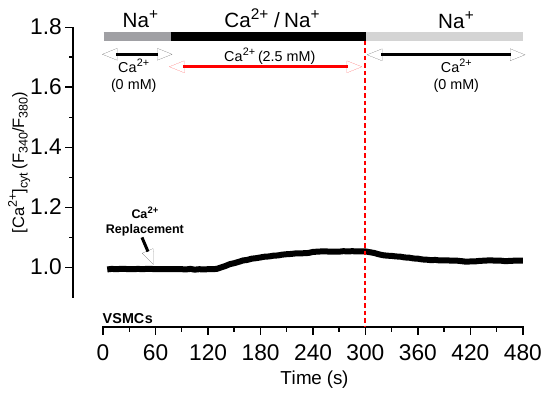

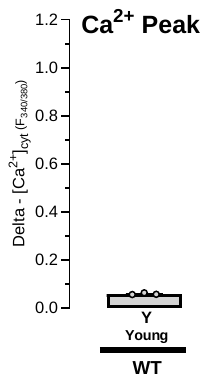

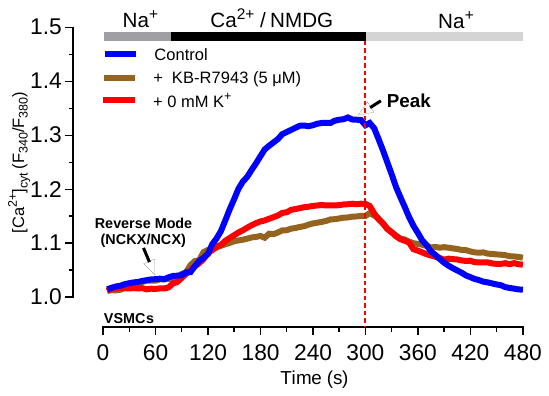

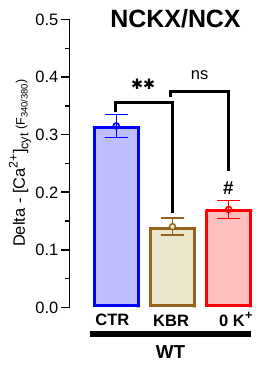

**A**

**B**

**C**

**D**

**Figure S6.** Chemical and pharmacological validation of NCKX/NCX activity in Ca^2+^ influx reverse mode in primary VSMCs from young (Y, 12-15 weeks) female wild-type (WT) mice loaded with the Ca²⁺ probe Fura-2-AM. (**A**)**,** effects of the Ca^2+^ replacement on Ca^2+^ transients in in VSMCs from young WT mice. A free Ca^2+^-solution perfused for 60s was replaced by normal Ca^2+^ solution containing 2.5 mM of CaCl_2_. (**B**), delta ∆ peak parameter was calculated. (**C**)**,** VSMCs showing NCKX/NCX activity in Ca^2+^ influx reverse mode. This mode was elicited by the replacement of Na^+^ with N-Methyl-D-glucamine (NMDG) and the addition of Ca^2+^ in the presence or absence of K^+^ solution (4.5 mM KCl). To distinguish NCX from NCKX function in Ca^2+^ influx mode, we halted NCKX activity by replacing the bath solutions with a K^+^-free solution evoking a decrease in Ca^2+^ transients (red trace). Ca^2+^ transients were measured in the presence of the NCX inhibitor KBR7943 (30 min, 5 μM; amber trace). **(D)**, quantification of the Ca^2+^ influx peak based on data from C. Data represent the mean ± SEM of ≥34 cells analyzed by one-way ANOVA followed by Tukey’s multiple comparison post-hoc test. **P < 0.01 control vs KBR group; #P < 0.05 control vs 0K^+^ group; n.s, nonsignificant.

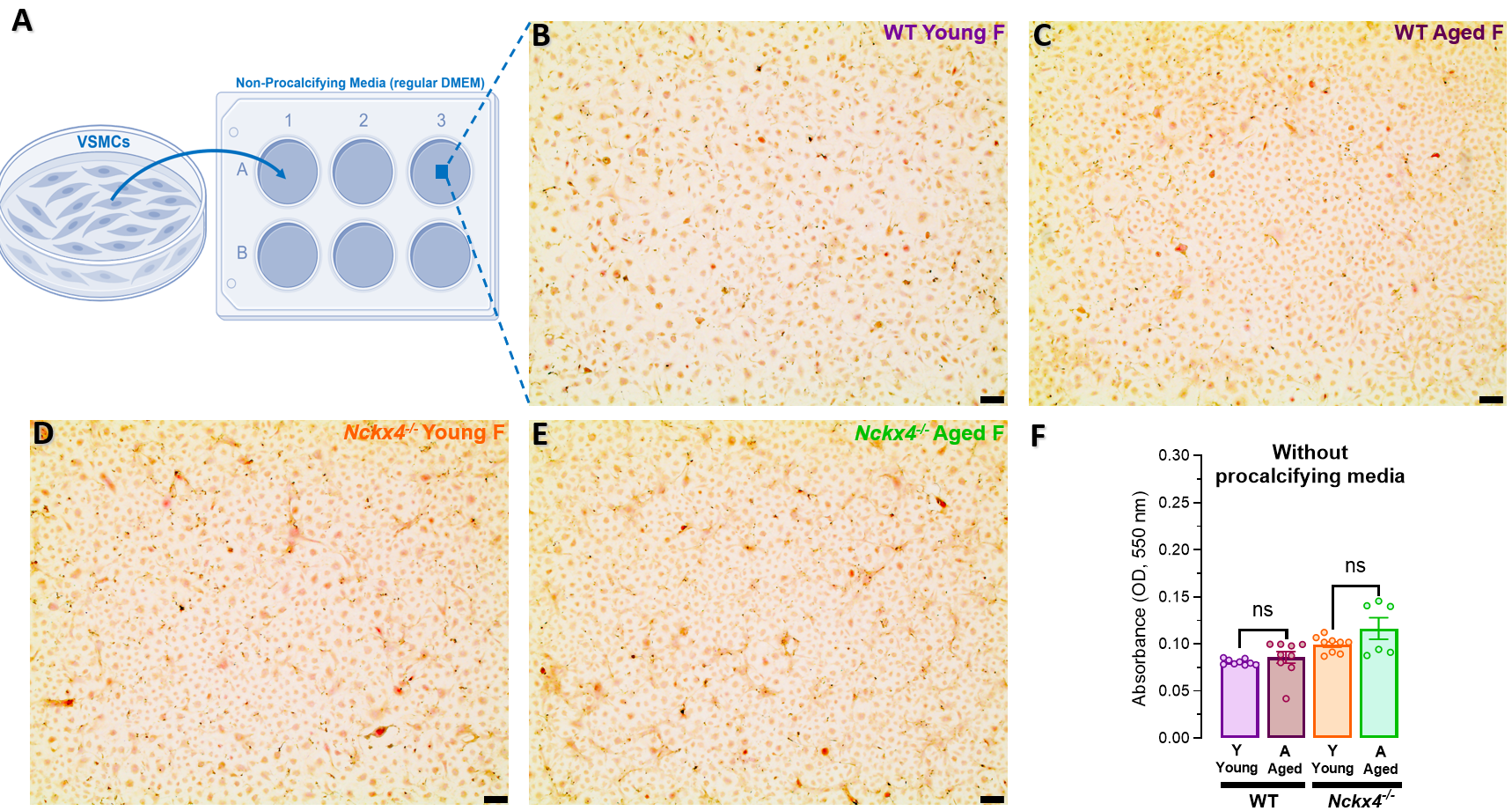

**Figure S7.** Alizarin Red-S staining in primary VSMCs from young (Y, 12-15 weeks) and aged (A, 72-78 weeks) male and female *Nckx4* knockout (*Nckx4^-/-^*) and wild-type (WT) mice. The VSMCs were culture in a non-procalcifying control media (high glucose regular DMEM media) for 16 days. (**A-E**), schematic and representative images of control VSMCs after 16-days using a non-procalcifying control media. VSMCs from young (Y, 12-15 weeks) and aged (A, 72-78 weeks) female wild-type (WT) and *Nckx4^-/-^* mice loaded were stained with Alizarin Red-S (2%, w/v), and dissolved by hexadecyl pyridinium (10%, w/v). Data represent the mean ± SEM of ≥3 experiments, and each individual dot in the histograms represent an independent sample from 6-well-plate stained with Alizarin Red-S. Data were analyzed by one-way ANOVA followed by Tukey’s multiple comparison post-hoc test. Significance was accepted at: *P<0.05; ***P<0.001; ^#^P< WT young group vs. *Nckx4^-/-^* young group.

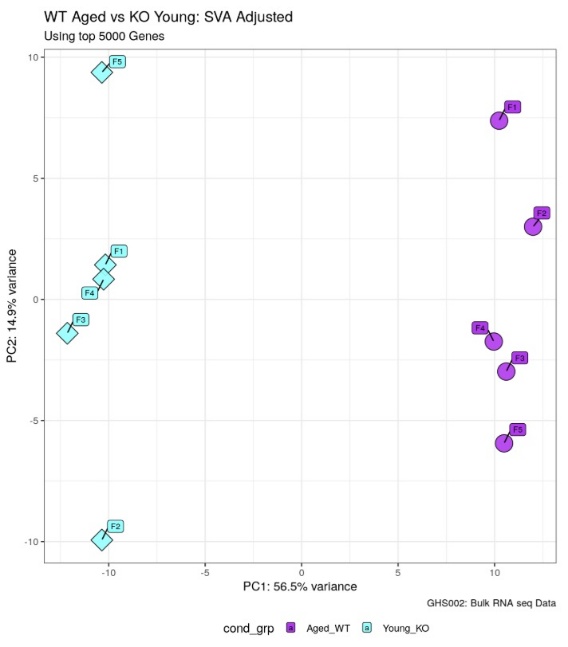

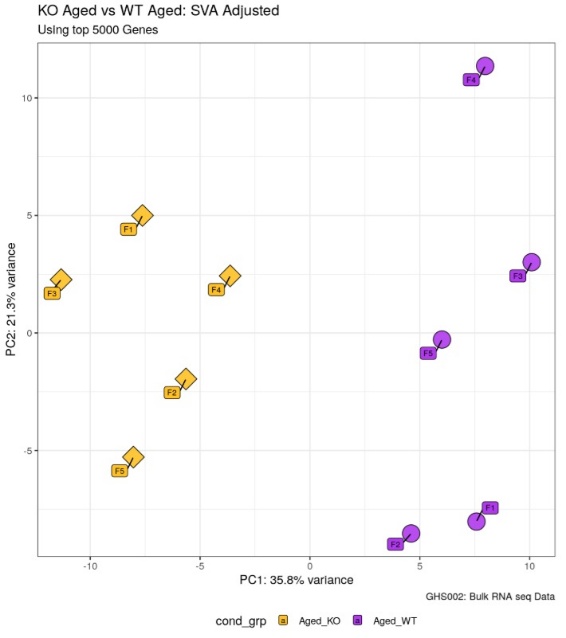

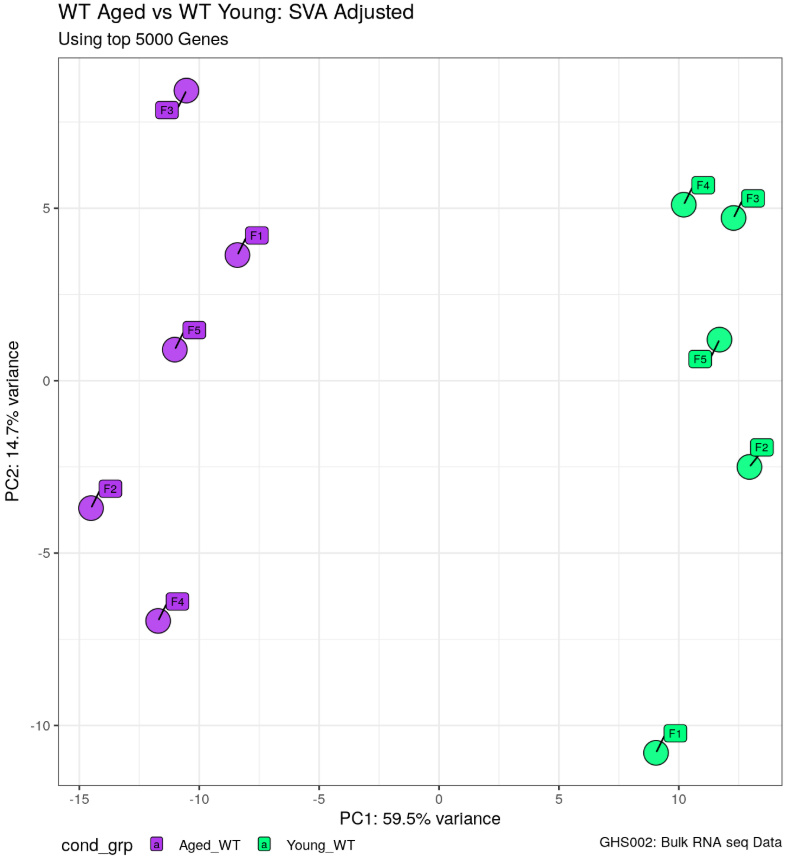

**A**

**B**

**C**

**Figure S8.** Principal Component Analysis (PCA) of bulk RNA-seq data of aortic tissue from young (Y, 12-15 weeks) and aged (A, 72-78 weeks) female wild-type (WT) and *Nckx4^-/-^* mice. PCA was conducted on variance-stabilized gene expression values for all four experimental groups. **(A)**, WT aged vs WT young. **(B)**, WT aged vs KO (*Nckx4^-/-^*) young. **(C)**, KO (*Nckx4^-/-^*) aged vs WT aged. Each point represents an individual biological replicate, total of 5 different samples.

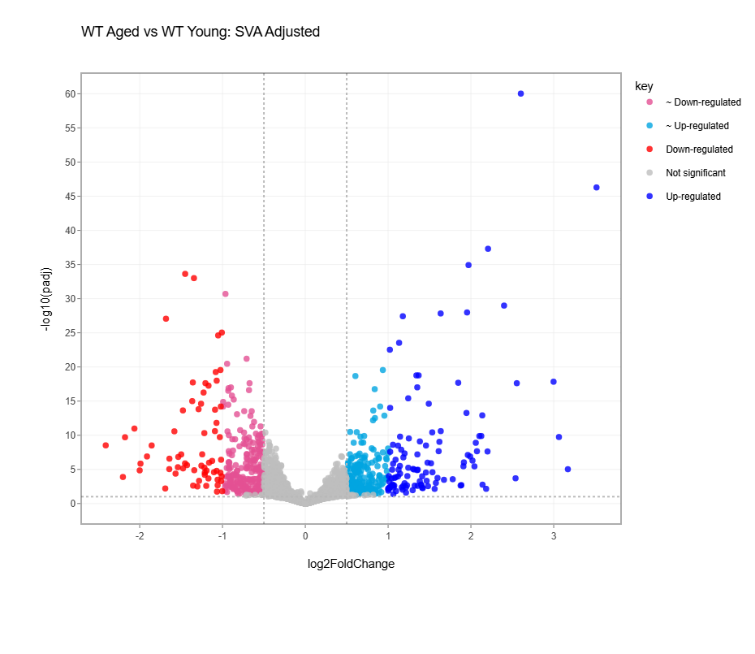

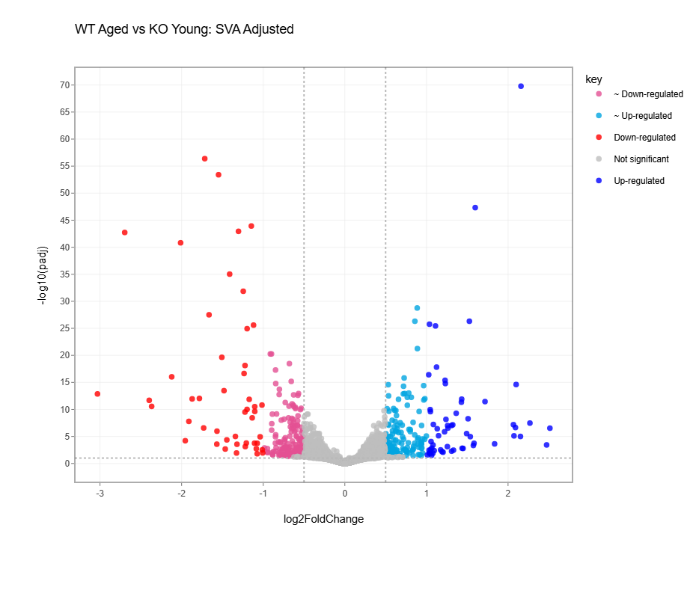

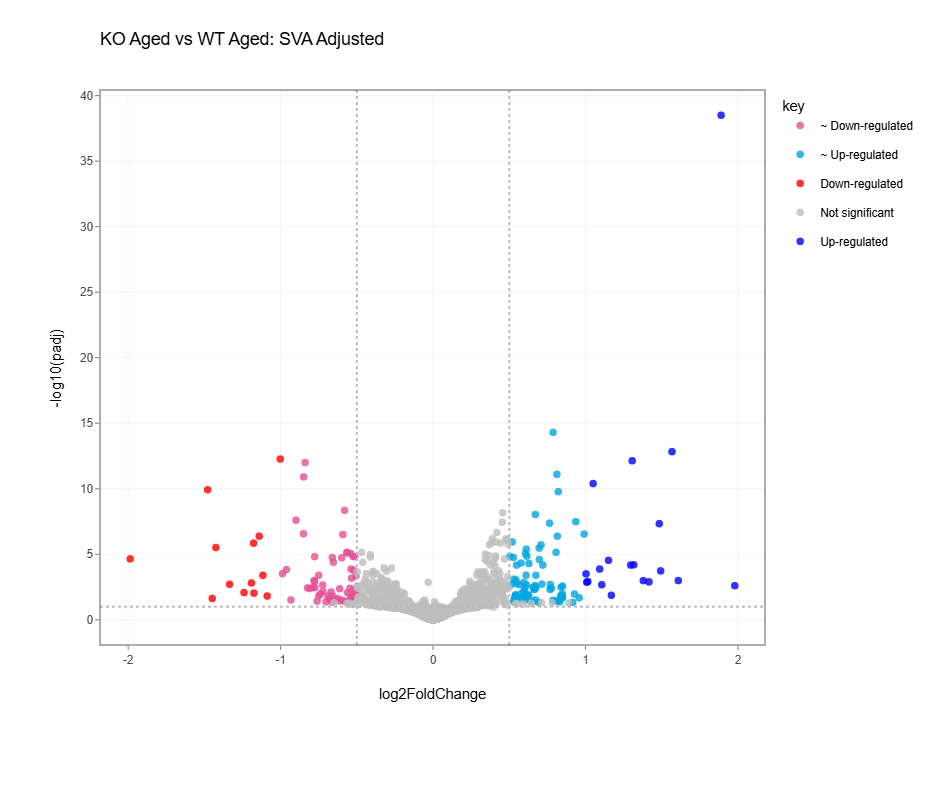

**A**

**B**

**C**

**Figure S9.** Volcano plot of differential gene expression of bulk RNA-seq data of aortic tissue from young (Y, 12-15 weeks) and aged (A, 72-78 weeks) female wild-type (WT) and *Nckx4^-/-^* mice. The volcano plot displays the distribution of all analyzed genes based on their differential expression between experimental conditions. **(A)**, WT aged vs WT young. **(B)**, WT aged vs KO (*Nckx4^-/-^*) young. **(C)**, KO (*Nckx4^-/-^*) aged vs WT aged. Each point represents an individual gene significantly upregulated (log₂FC > threshold, padj < 0.05) colored in red (light and dark), while significantly downregulated genes (log₂FC < threshold, padj < 0.05) in blue (light and dark). Genes with no significant change are plotted in gray.

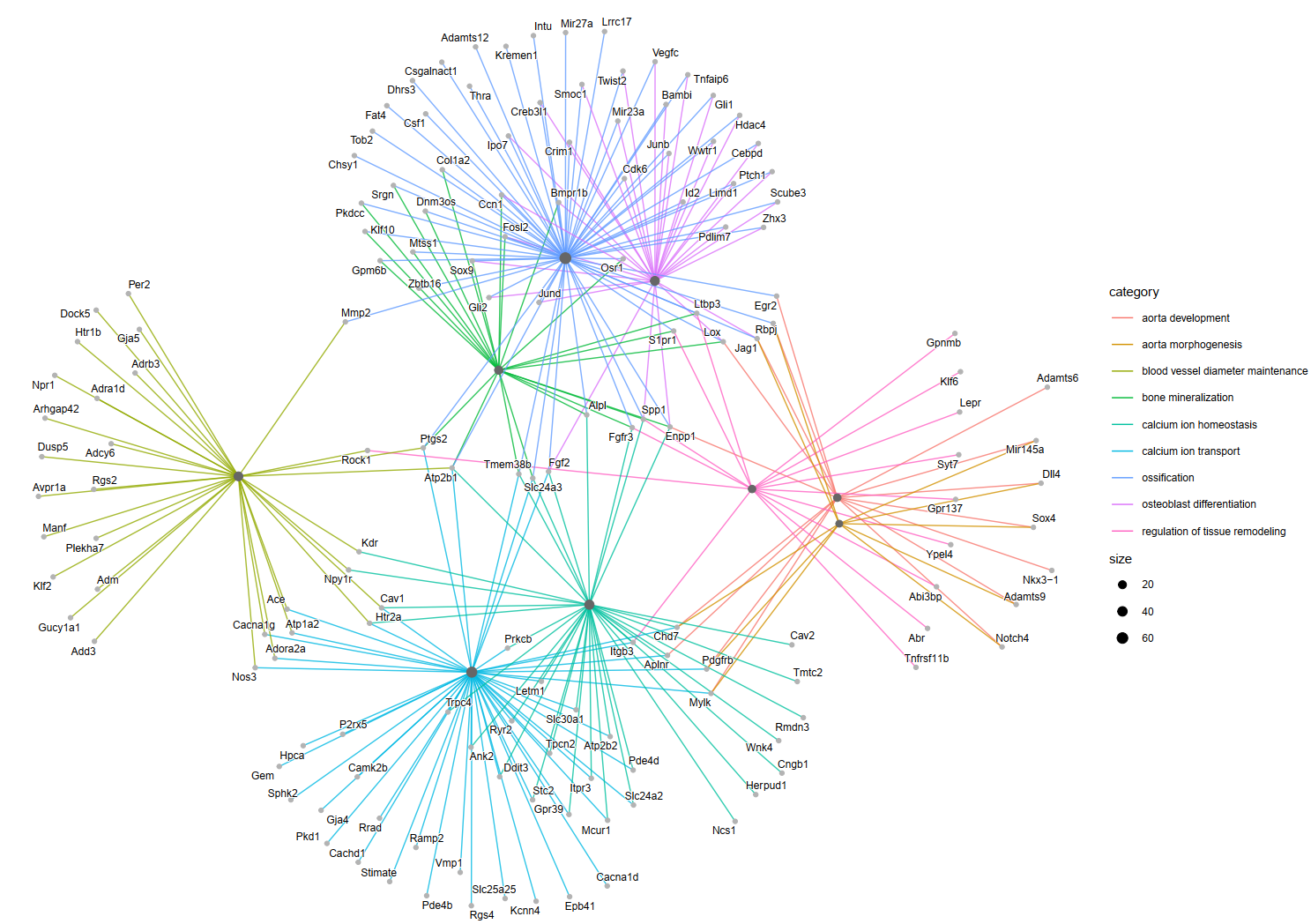

**A**

**B**

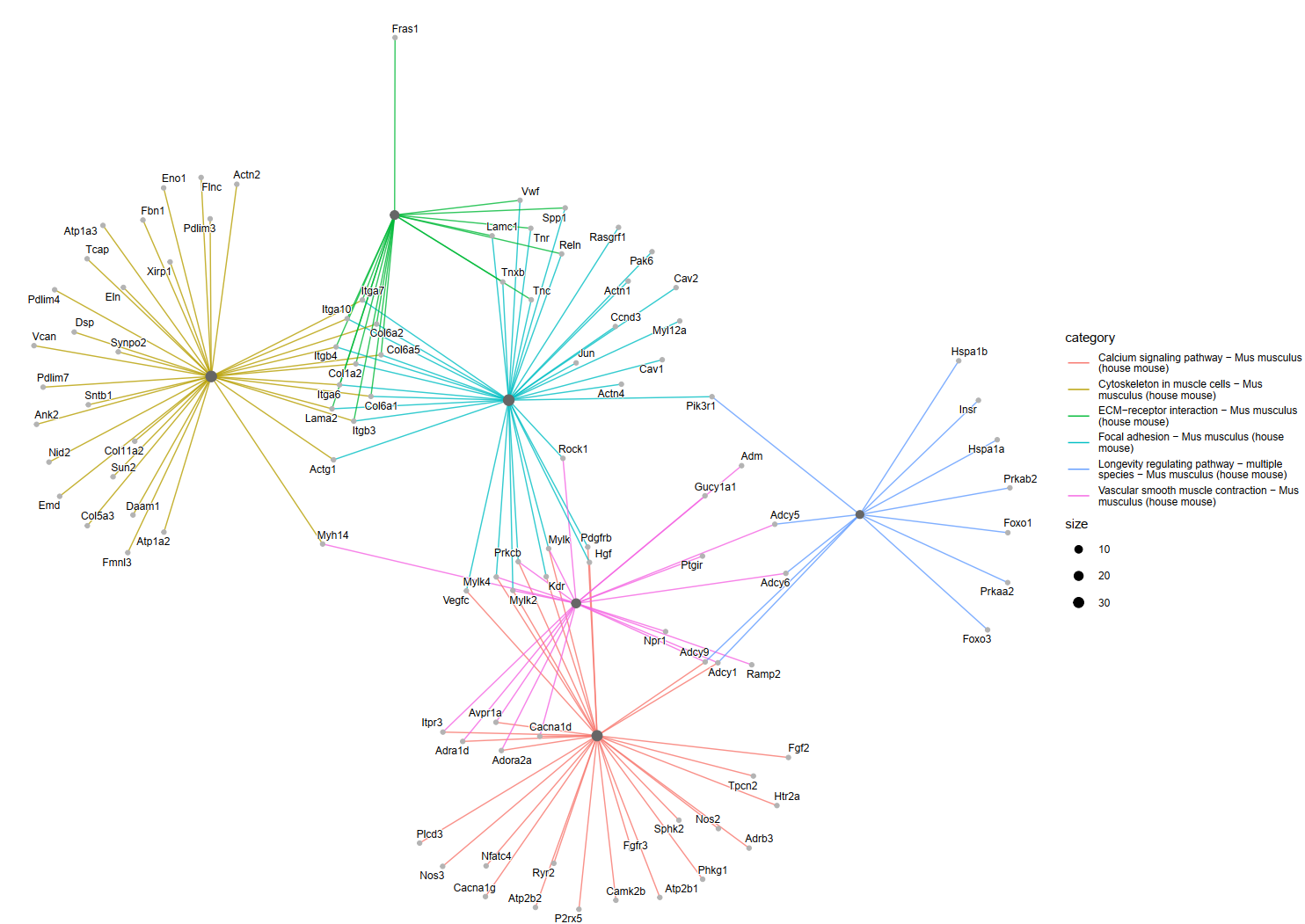

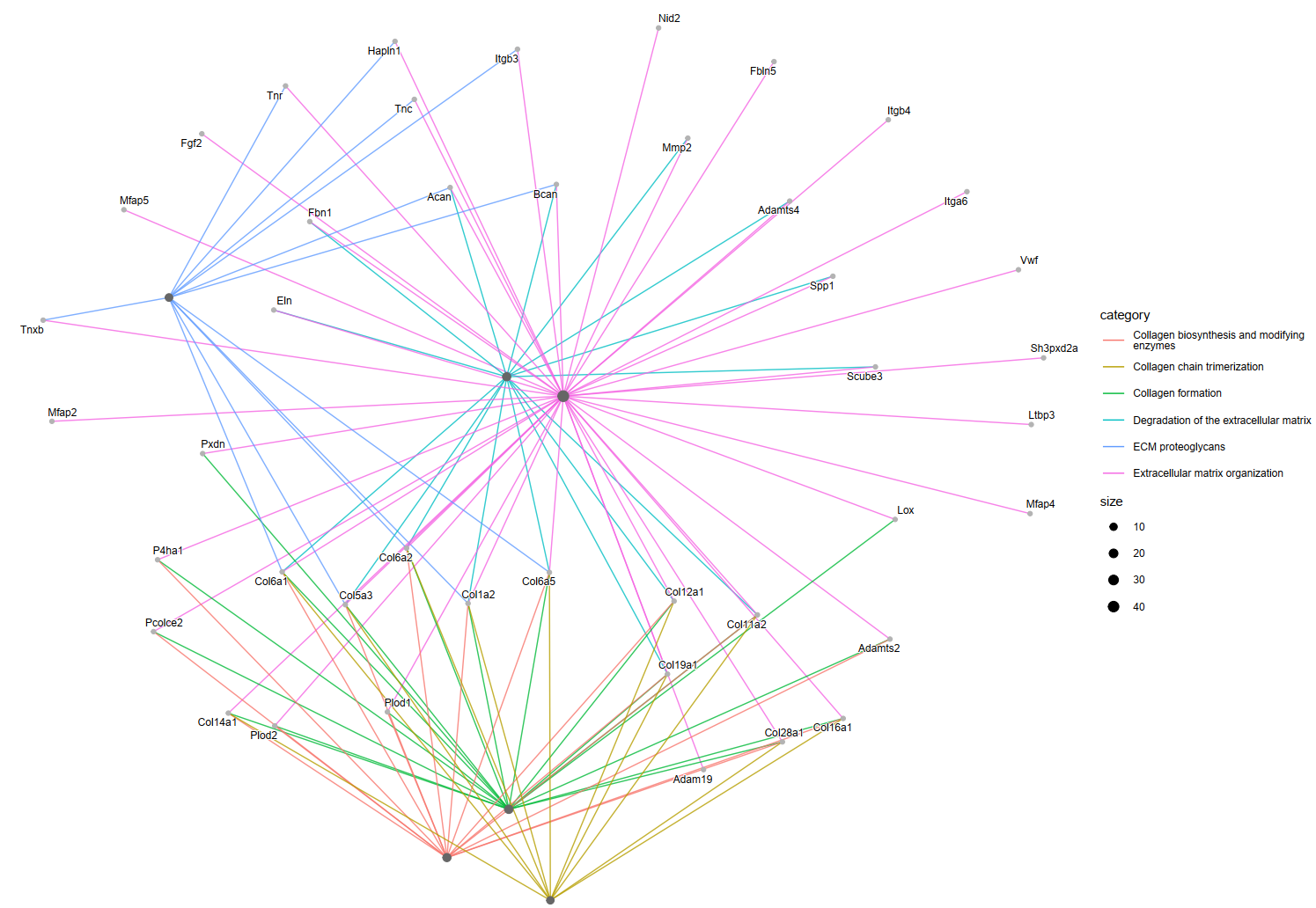

**C**

**Figure S10.** Differential expression analysis of pathways identified genes with significantly different expression using gene regulatory network representation. GO BP **(A)**, KEGG **(B)** and Reactome **(C)** in aortic tissue from young (Y, 12-15 weeks) *Nckx4^-/-^* mice when compared with aged (A, 72-78 weeks) female wild-type (WT) mice.
